# Mitochondrial introgression in North American red-backed voles is facilitated by co-introgression at nuclear-encoded mitochondrial genes

**DOI:** 10.64898/2026.07.22.740054

**Authors:** Ben J. Wiens, Jocelyn P. Colella

## Abstract

Mitochondrial genomes encode for the proteins and RNAs that serve as the basis for aerobic respiration and energy production. Yet mitochondria depend on the nuclear genome for hundreds of additional genes whose products engage in core metabolic processes (N-mt genes). Despite strong selective pressure for coevolved mitonuclear interactions, there are numerous examples of mitochondrial introgression across species barriers. Mitonuclear co-introgression, a process whereby alleles at N-mt genes move across species boundaries in concert with mitochondrial genomes, has been suggested as a mechanism whereby species could capture heterospecific mitochondria while avoiding mitonuclear incompatibilities, but evidence for this phenomenon is sparse. We test for evidence of mitonuclear co-introgression in two discordant populations of North American red-backed voles (*Clethrionomys gapperi* nuclear genomes, *C. rutilus* mitochondrial genomes) using whole genome resequencing. We find that N-mt genes in both populations are significantly enriched for *C. rutilus* ancestry, with evidence of co-introgression at eighteen N-mt genes. Notably, two N-mt genes directly associated with mitochondrial translation are fixed or nearly fixed for *C. rutilus* ancestry in both discordant populations and analyses of genetic variation at these genes suggest recent selective sweeps. We pair these findings with mitochondrial phylogenies, recent demographic histories, and recombination maps, which support a scenario of ongoing introgression in British Columbia but cessation of gene flow in Southeast Alaska. Together, our results show that mitochondrial introgression in North American red-backed voles is adaptive and that mitonuclear incompatibilities are avoided through mitonuclear co-introgression.

## 1 | Introduction

Mitochondria provide the key to aerobic respiration in eukaryotic organisms, and as the result of ancient symbiosis, retain a genome separate from the cell nucleus (Hill, 2015; Spinelli & Haigis, 2018). Over time, mitochondrial genes have been lost or incorporated into the nuclear genome, but mitochondrial genomes in almost all animals still contain 13 protein coding regions, 22 tRNAs, 2 rRNAs, and a small regulatory region (Adams & Palmer, 2003; Chinnery & Hudson, 2013). Those genes encode for the major subunits of the electron transport chain, which enable oxidative phosphorylation, while mt-tRNAs and and mt-rRNAs facilitate mitochondrial translation (Chinnery & Hudson, 2013; Diodato et al., 2014). While directly responsible for the conversion of organic compounds into usable cellular energy, mitochondria are also essential for a number of other cellular processes, including biosynthesis, thermogenesis, cell signaling, stress responses, and immunity (Chouchani et al., 2019; Lajbner et al., 2018; Marques et al., 2024; Spinelli & Haigis, 2018; Wallace, 2016).

Given their involvement in multiple fundamental molecular pathways, mitochondrial products must interact with products of the nuclear genome (Hill, 2015; Rath et al., 2021). Genes in the nuclear genome that facilitate mitochondrial processes, either directly or indirectly, are referred to as nuclear-encoded mitochondrial genes (N-mt genes); in mammals, ∼1,140 N-mt genes have been described (Chinnery & Hudson, 2013; Rath et al., 2021). Within a species, selection acts to maintain compatible gene interactions across the mitochondrial and nuclear genomes, despite their distinctly different modes of inheritance (Hill, 2015; Rand et al., 2004). However, mitonuclear incompatibilities can arise when mitochondrial genomes are placed against a novel background of nuclear genotypes, which can happen as a result of hybridization (Hill et al., 2019; Sloan et al., 2023). In this context, mitonuclear incompatibilities are a form of Dobzhansky-Muller incompatibilities (DMIs), which arise when independently derived alleles at two or more loci no longer interact properly in hybrids (Burton & Barreto, 2012). Since mitochondrial genomes generally experience higher substitution rates than in the nuclear genome and are inherited matrilineally, it has been suggested that the disruption of mitonuclear interactions may be one of the most common forms of hybrid incompatibilities (Allio et al., 2017; Ballard & Whitlock, 2004; Hill et al., 2019). Indeed, there are many examples of severe mitonuclear incompatibilities between closely related species (Lee et al., 2008; Meiklejohn et al., 2013; Moran et al., 2024; Niehuis et al., 2008) and even between populations of single species (Chang et al., 2016; Lima et al., 2019; Ma et al., 2016).

Despite theoretical and empirical evidence that mitonuclear mismatch can give rise to genetic incompatibilities, mitochondrial capture is frequently observed in nature (Toews & Brelsford, 2012). A variety of verbal models are often invoked to explain this phenomenon, most of which rely on demographic- or sex-based biases in mating or dispersal and the fact that mitochondrial genomes are inherited matrilineally (Bonnet et al., 2017; Toews & Brelsford, 2012). Yet, when simulated, those models fail to produce the signature of large-scale mitochondrial introgression and minimal nuclear introgression that they attempt to explain (Bonnet et al., 2017). Instead, those simulations found support for adaptive mitochondrial introgression as a major driver of mitochondrial capture. Numerous empirical studies further support the notion that introgression of heterospecific mitochondrial genomes can provide an adaptive advantage, with evidence from brook char (Doiron et al., 2002), frogs (Plötner et al., 2008), buntings (Irwin et al., 2009), robins (Morales et al., 2018), siskins (Beckman et al., 2018), bank voles (Boratyński et al., 2014), goats (Ropiquet & Hassanin, 2006), bears (Hailer et al., 2012), and hares (Melo-Ferreira et al., 2005; Melo-Ferreira et al., 2014), for example. While introgressed mitochondria could provide a selective advantage through numerous pathways, a commonly proposed mechanism is by increased cold tolerance through elevated thermogenesis (Hill, 2019; Lajbner et al., 2018). Aligned with that hypothesis, mitochondrial genes are known to be involved in local adaptation to thermal gradients in mammals (Fontanillas et al., 2005; Grover-Thomas et al., 2026), birds (Morales et al., 2015), and fish (Silva et al., 2014).

It is easiest for mitochondrial introgression to occur when there are no mitonuclear incompatibilities between the hybridizing species, but it is not limited to this scenario (Hill, 2019; Sloan et al., 2023). For example, if a heterospecific mitochondrial genome provides a selective advantage that outweighs the disadvantage of accompanying mitonuclear incompatibilities, mitochondrial introgression could still occur. Alternatively, mitonuclear incompatibilities could be mitigated through co-introgression of heterospecific nuclear ancestry at the N-mt genes involved in the incompatibility (Beck et al., 2015; Sloan et al., 2023). Distinguishing between these scenarios should be possible by analyzing variation across the nuclear and mitochondrial genomes in populations that exhibit mitonuclear mismatch. If there are no mitonuclear incompatibilities, or if they are offset by the advantage of heterospecific mitochondria, then ancestry patterns at N-mt genes should be the same as seen across the rest of the nuclear genome. Alternatively, if mitonuclear incompatibilities exist but are avoided through mitonuclear co-introgression, then N-mt genes should be enriched for heterospecific ancestry.

We test for empirical evidence of mitonuclear co-introgression in North American red-backed voles (*Clethrionomys*). Previously, we used reduced-representation sequencing to document mitochondrial introgression from the northern red-backed vole (*C. rutilus*) into populations of the southern red-backed vole (*C. gapperi*) in zones of contact in Southeast Alaska and British Columbia (Wiens & Colella, 2025). Unidirectional introgression of *C. rutilus* mitochondria, which evolved in Beringia during Pleistocene glacial cycles (Kohli et al., 2015), into populations with *C. gapperi* nuclear ancestry, which evolved south of the Pleistocene ice sheets in North America (Runck & Cook, 2005), suggests that *C. rutilus* mitochondria may provide a selective advantage for *C. gapperi* populations as they expand north into colder climates. If mitochondrial introgression from *C. rutilus* to *C. gapperi* is adaptive, but mitonuclear incompatibilities exist, we expect to find increased *C. rutilus* ancestry at N-mt genes in discordant populations. In this study, we characterize multiple aspects of genomic variation in two red-backed vole populations that have experienced mitochondrial introgression. First, we build recombination maps, infer recent demographic histories for each species, and compare histories of contact and admixture in Southeast Alaska and British Columbia. We then reconstruct local ancestry across the nuclear genome, test for evidence of mitonuclear co-introgression, and infer patterns of selection at N-mt genes. Together, our results shed light on the mechanisms that lead to mitochondrial introgression across species boundaries.

## 2 | Results

### Genome-wide ancestry patterns and history of secondary contact

We characterized patterns of nuclear and mitochondrial ancestry across Southeast Alaska (SEAK) and British Columbia (BC) with whole genome resequencing data for 72 red-backed voles (Figure 1). We focused our sampling on the center of the contact zone in each region, but included 21 *C. rutilus* and 13 *C. gapperi* from outside the contact zone in British Columbia to use as reference populations for each species. To assess hybrid ancestry in the nuclear genome, we developed a dataset of 234,874 biallelic single nucleotide polymorphisms (SNPs). We then identified 25,375 SNPs with fixed differences between the reference populations, which we used as ancestry-informative markers (AIMs) to calculate hybrid index and interclass heterozygosity with *triangulaR* (Wiens et al., 2025). We also assembled whole mitogenomes from our resequencing data and used *iqtree* (Minh et al., 2020) to build a mitochondrial phylogeny (Figure 2). Through the combination of nuclear and mitochondrial data, we identify five ancestry groups across the landscape, which match previous findings based on RADseq data (Wiens & Colella, 2025). In Southeast Alaska, red-backed voles north of the LeConte Bay are *C. rutilus* and red-backed voles on southern, interior islands of the Alexander Archipelago are *C. gapperi*. On the coastal mainland south of the LeConte Bay, there is a discordant population of red-backed voles with *C. rutilus* mitochondrial genomes and majority *C. gapperi* nuclear ancestry (98.6% *C. gapperi* ancestry on average). There is also a discordant population with *C. rutilus* mitochondrial ancestry and majority *C. gapperi* nuclear ancestry (92.2% *C. gapperi* on average) in British Columbia. Less than 10 km to the north, populations turnover to full *C. rutilus* ancestry, while mitonuclear discordance extends for at least 100 km to the south (Figure 1; Wiens & Colella, 2025).

**Figure 1.**
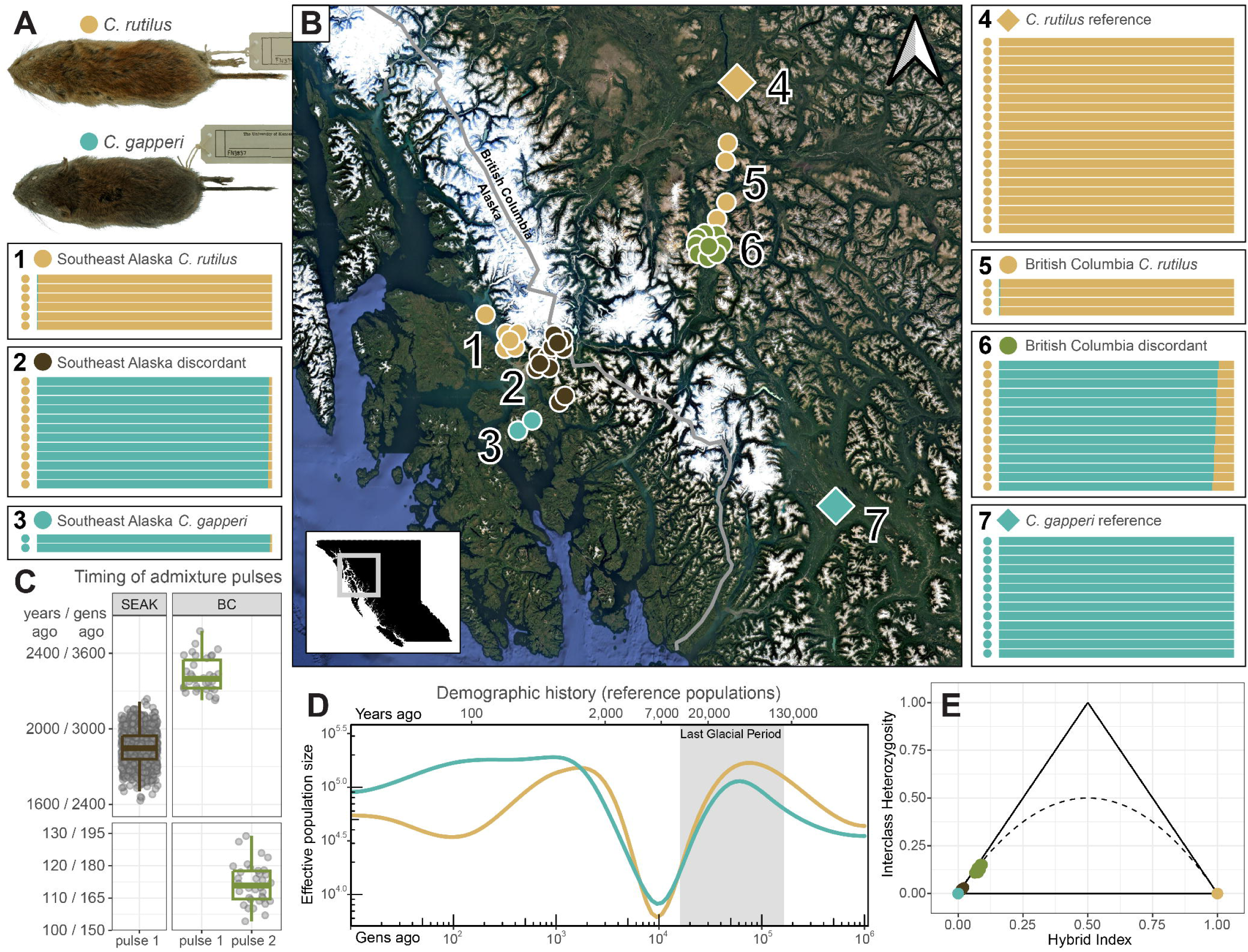
**(A)** Representative specimens of *Clethrionomys rutilus* and *C. gapperi*. **(B)** Locations of sampled red-backed voles from Southeast Alaska (SEAK) and British Columbia (BC). Colors on the map indicate mitonuclear ancestry (*C. gapperi* = teal; *C. rutilus* = gold; SEAK discordant = brown; BC discordant = green) and numbers on the map identify each sampled population and correspond to the panels to the left and right of the map. Diamonds represent the reference populations of each species and circles represent individual voles. Each numbered panel contains a row for each individual vole sampled from that population. Each row consists of a colored circle on the left indicating the mitochondrial species ancestry and a bar on the right indicating average ancestry proportions across the nuclear genome, calculated using 25,375 AIMs in *triangulaR*. **(C)** Timing of admixture pulses for discordant populations in Southeast Alaska (brown) and British Columbia (green). For each, only the best fitting model is shown. Grey points represent 50 (2 pulse model) or 1,000 (1 pulse model) bootstrap estimates of the timing of an admixture pulse in terms of generations (gens) before present, and are summarized by a boxplot. Generations were converted to years by assuming 1.5 generations per year. **(D)** Inferred demographic histories based on the reference population of each species (*C. gapperi* = teal; *C. rutilus* = gold). Grey box indicates the peak of the Last Glacial Period (10,700 to 110,000 years before present). **(E)** Triangle plot built based on 25,375 AIMs. Only the reference populations and the two discordant populations are shown.

**Figure 2.**
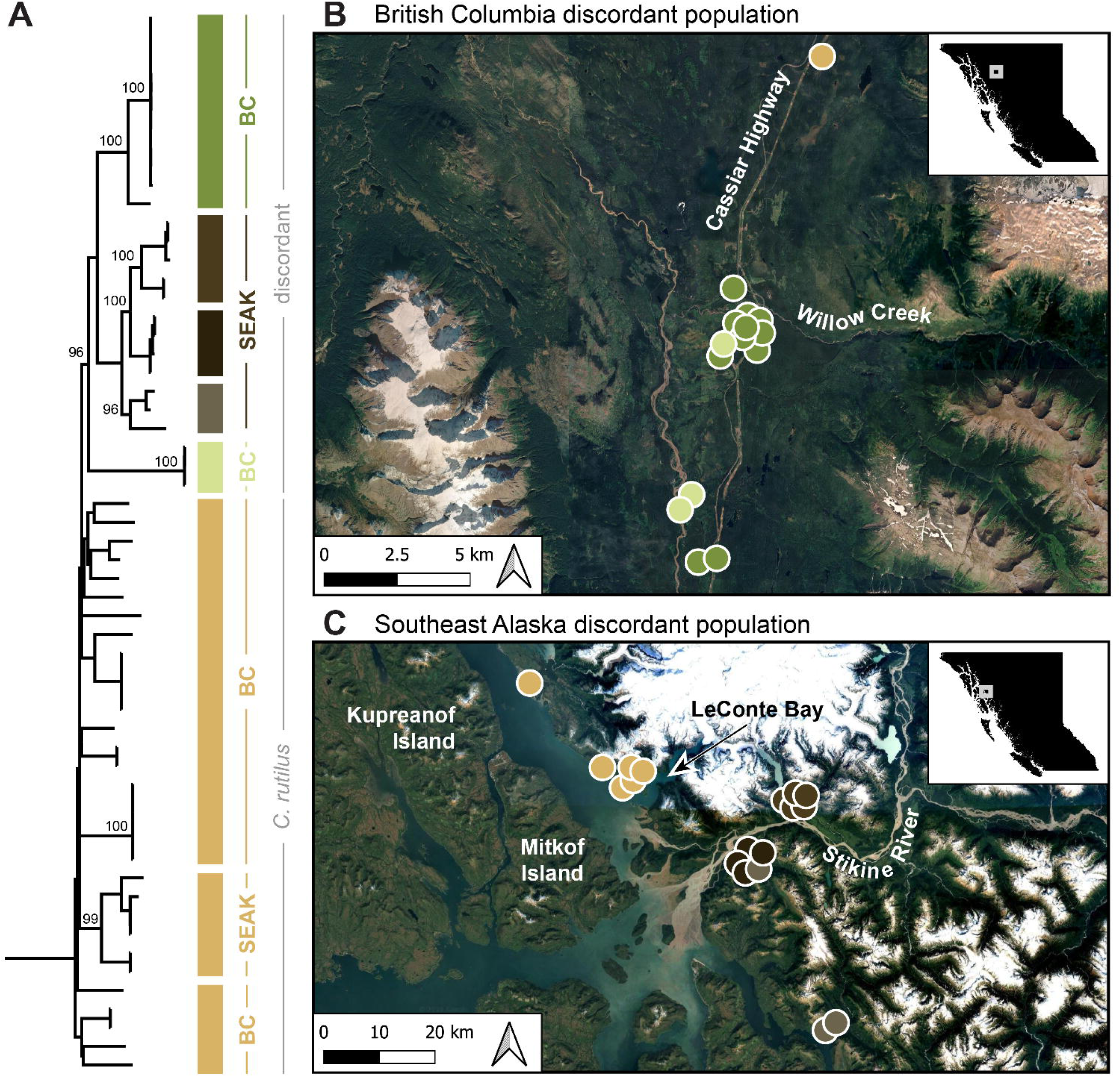
**(A)** Maximum likelihood mitogenome phylogeny for voles with *C. rutilus* mitochondria. Branches with >95% bootstrap support are labeled. **(B)** Map of the center of the contact zone in British Columbia. Each point represents an individual vole colored by mitochondrial clade. The nearest full *C. rutilus* individual is shown at the top of the map, all other *C. rutilus* from British Columbia seen in the mitogenome phylogeny were sampled farther north. **(C)** Map of the center of the contact zone in Southeast Alaska.

We took two approaches to understand the history of hybridization between *C. rutilus* and *C. gapperi*. First, we inferred demographic histories for each species using a sequentially Markov coalescent approach, implemented in *smc++* (Terhorst et al., 2017). For both species, we infer a rapid decline in effective population size following the last glacial maximum (∼20 kya), consistent with leading edge effects expected from a rapid expansion out of glacial refugia, followed by a quick return to pre-expansion effective population sizes (Figure 1D; Hewitt, 1996). Second, we used a hidden Markov model, as implemented in *Ancestry_HMM* (Medina et al., 2018), to simultaneously infer local ancestry and the timing of contact between *C. rutilus* and *C. gapperi* in each geographic region based on the distribution of observed ancestry tract lengths in each discordant population. For each discordant population, we compare models that assume either one or two pulses of admixture and estimate when those pulses occurred, bounding the possible pulses to within the last 1,000,000 generations (Figure 1C). In Southeast Alaska, admixture is estimated at 2,850 generations ago (1,000/1,000 bootstrap support) by the one-pulse model and 999,998 and 2,798 generations ago (50/50) by the two-pulse model. In British Columbia, admixture is estimated at 1,003 generations ago (1,000/1,000) by the one-pulse model, while the two-pulse model is split between two estimates: pulses at 3,428 and 171 generations ago (34/50) and pulses at 1,000,000 and 981 generations ago (16/50). It is difficult to select between one-and two-pulse admixture models given that the additional parameters of the two-pulse model almost always provide a better fit to the data. To select between models, one heuristic suggested by the authors of *Ancestry_HMM* is that if the initial admixture pulse in the two-pulse model is estimated at the maximum bound of the time interval, then the one-pulse model is likely a better fit (Medina et al., 2018). Following that suggestion, our data support a single pulse of admixture in Southeast Alaska around 2,850 generations ago and multiple admixture pulses in British Columbia, around 3,428 and 171 generations ago.

### Biased nuclear introgression at N-mt genes

Mitochondrial genes interact either directly or indirectly with ∼1,140 N-mt genes, giving rise to the possibility of mitonuclear incompatibilities when mitochondrial and nuclear genes are mismatched following admixture events (Hill et al., 2019; Rath et al., 2021). When mitochondrial genomes do move across species boundaries, one way that mitonuclear incompatibilities can be avoided is through co-introgression of N-mt genes (Sloan et al., 2017). Recombination may facilitate biased introgression of N-mt genes by breaking apart unfit haplotypes and allowing selection to act on N-mt genes in novel genomic backgrounds (Sloan et al., 2017). The small amount of *C. rutilus* nuclear ancestry in each discordant population (1.4% in SEAK, 7.8% in BC) suggests the possibility of mitonuclear co-introgression. Alternatively, *C. rutilus* nuclear ancestry in the discordant populations could simply be remnants from past admixture events and randomly dispersed across the genome.

We characterized patterns of genomic variation in each discordant population by pairing local ancestry maps with estimates of recombination rate variation, nucleotide diversity (π), and genetic differentiation (F_ST_, D_XY_) between the reference populations of each species (Figure 3). We find high genetic differentiation between *C. rutilus* and *C. gapperi* (average sliding window F_ST_ = 0.72, D_XY_ = 0.0068), while nucleotide diversity is similar within each species (*C. rutilus* π = 0.0018; *C. gapperi* π = 0.0016). Recombination rates in sliding windows across the genomes are similar (*C. rutilus* r = 3.20 × 10^-9^; *C. gapperi* r = 3.87 × 10^-9^), correlated (Spearman’s ρ = 0.44, *P* < 2.2 × 10^-16^), and match expectations based on karyotypes. Both species have 27 autosomes, all of which are acrocentric except for chromosome 27, which is metacentric (Nadler et al., 1976; Rausch & Rausch, 1975). We find evidence for reduced recombination near the center of chromosome 27, likely corresponding to the centromere, while for the other 26 chromosomes there is no major reduction in recombination near the chromosome center (Figure 3, Figure S1).

**Figure 3.**
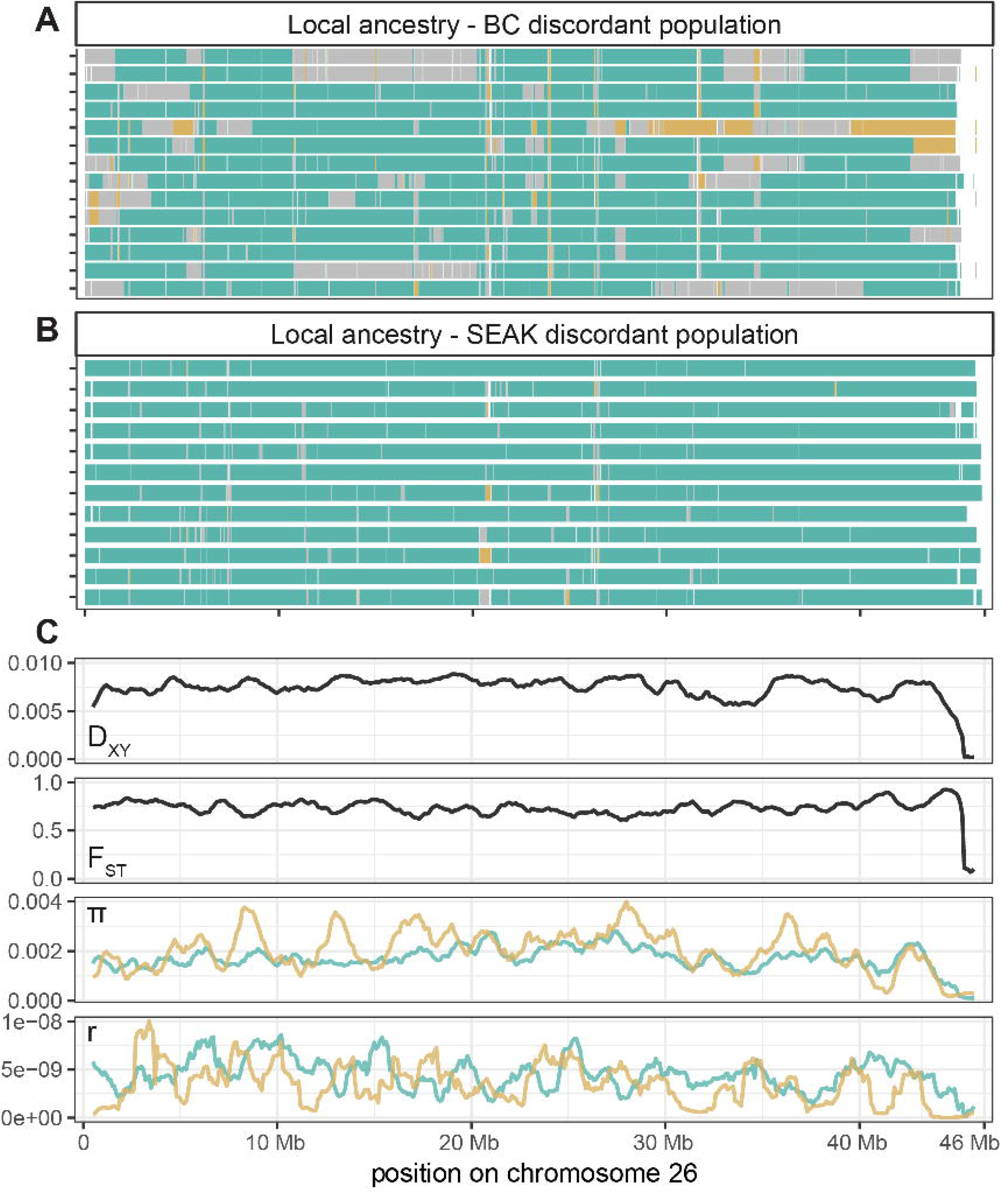
Local ancestry along chromosome 26 for each individual in the **(A)** British Columbia (BC) discordant population and **(B)** Southeast Alaska (SEAK) discordant population. Each individual is represented by a single horizontal bar. Species ancestry at each position along the chromosome is indicated by color: *C. gapperi* = teal; *C. rutilus* = gold; heterozygous = grey. **(C)** Patterns of genetic variation along chromosome 26 for the reference population of each species. Genetic differentiation (F_ST_, D_XY_) between reference *C. gapperi* and reference *C. rutilus* is summarized in 1 Mb windows, slid across the chromosome in 100 kb intervals. Nucleotide diversity (π) and recombination rate (r) in the same windows are shown for each reference population (*C. gapperi* = teal; *C. rutilus* = gold).

To test for evidence of mitonuclear co-introgression we calculated the average proportion of *C. rutilus* ancestry in N-mt genes and in equally-sized genomic windows for each discordant population. Of 1,140 N-mt genes in Mouse MitoCarta3.0 (Rath et al., 2021), 1,027 are annotated in the *C. rutilus* reference genome. Some of the genes have multiple copies in the *C. rutilus* reference genome, for a total of 1,124 N-mt loci. For the discordant populations, we find that *C. rutilus* ancestry at N-mt genes is higher than expected given the distribution of *C. rutilus* ancestry across the genome (Kolmogorov-Smirnov test: SEAK mean ancestry at N-mt genes = 0.025, genomic windows = 0.016, *P* = 0.033; BC mean ancestry at N-mt genes = 0.100, genomic windows = 0.079, *P* = 3.015 × 10^-4^). That pattern is primarily driven by a subset of N-mt genes with high amounts of *C. rutilus* ancestry (Figure 4A-B). To provide additional support for excess *C. rutilus* ancestry at N-mt genes, we then tested the probability that a random sample of 1,124 genomic windows could contain the average amount of *C. rutilus* ancestry observed at N-mt genes. For both discordant populations, the amount of *C. rutilus* ancestry at N-mt genes exceeds the upper 97.5% quantile of ancestry observed in 10,000 random samples of genomic windows (SEAK *P* = 1.7 × 10^-4^, BC *P* < 1.0 × 10^-6^), providing further support for co-introgression of *C. rutilus* mitochondria and N-mt genes (Figure 4C-D). Yet, we find no evidence for high recombination rates facilitating introgression of *C. rutilus* ancestry. Recombination rates at N-mt genes are low compared to other genomic windows (Kolmogorov-Smirnov test: SEAK *P* < 2.2 × 10^-16^, BC *P* < 2.2 × 10^-16^). Further, when only considering genomic windows without N-mt genes, we find that windows with high recombination rates are negatively associated with proportion of *C. rutilus* ancestry, although the association between those variables is very weak (SEAK *P* < 6.6 × 10^-9^, R^2^ = 4.4 × 10^-4^; BC *P* < 3.5 × 10^-8^, R^2^ = 4.0 × 10^-4^; Figure 4E-F).

**Figure 4.**
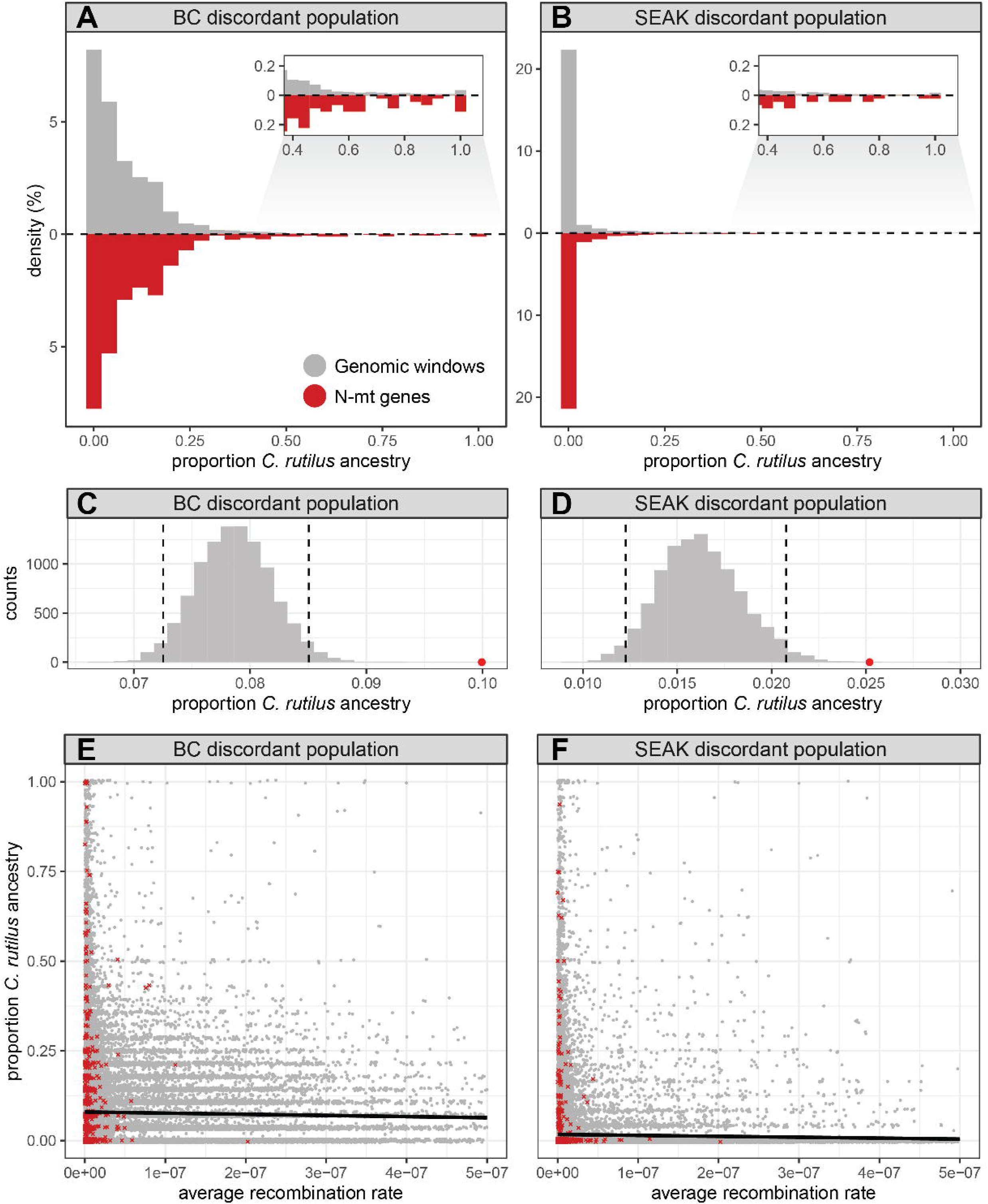
Proportion of *C. rutilus* nuclear ancestry in the discordant populations. **(A-B)** Relative frequency of *C. rutilus* nuclear ancestry in genomic windows (grey) and N-mt genes (red) for each discordant population. Insets show the upper tail of the distribution of *C. rutilus* ancestry proportions, from 0.4 to 1.0. **(C-D)** Histograms of the average proportion of *C. rutilus* nuclear ancestry in 10,000 random samples of 1,124 genomic windows. Black dotted lines indicate the lower 2.5% and upper 97.5% quantiles of the distributions. The red dot in each panel indicates the average proportion of *C. rutilus* ancestry at N-mt genes for each discordant population. **(E-F)** Relationship between recombination rate and the proportion of *C. rutilus* ancestry for genomic windows (grey) and N-mt genes (red). Linear regression reveals a significant, but slight negative relationship between recombination rate and proportion of *C. rutilus* ancestry in genomic windows in each discordant population (BC *P* < 3.5 × 10^-8^; SEAK *P* < 6.6 × 10^-9^).

### Positive selection for heterospecific ancestry at N-mt genes

After finding evidence for biased introgression of *C. rutilus* ancestry at N-mt genes overall, we conducted additional tests to identify specific N-mt genes with excess *C. rutilus* ancestry and signals of positive selection. We tested N-mt genes for higher proportions of *C. rutilus* ancestry than expected given the average amount of *C. rutilus* ancestry in each discordant population using a modified CAnD test (McHugh et al., 2016). After applying a Bonferroni correction, we find seventeen N-mt genes with significant excess *C. rutilus* ancestry in the BC discordant population and six in the SEAK discordant population, with five shared between them (Table 1). We visualize local ancestry in and around those genes for each discordant population, pairing local ancestry plots with sliding window estimates of F_ST_, D_XY_, recombination rate, and *Tajima’s D* (Figures 5-6, Figures S2-S37). We further tested those genes for evidence of positive selection by calculating *Tajima’s D* in each discordant population and the *C. rutilus* and *C. gapperi* reference populations. Of the eighteen identified N-mt genes, two show evidence of positive selection in both discordant populations (*Hemk1*: BC *D* = - 2.26, SEAK *D* = -2.48; *Kars*: BC *D* = -3.35, SEAK *D* = -2.06), four are under positive selection in *C. gapperi* (Table 1), and five are under positive selection in *C. rutilus* (Table 1). We then classified the mitochondrial pathways for each of the eighteen genes following MitoCarta3.0. Notably, twelve of these genes are involved in metabolism, with five associated with multiple metabolic pathways (Table 1). Further, two of the eighteen genes (*Hemk1* and *Kars*) are classified as part of the “mitochondrial central dogma”, defined by MitoCarta as mtDNA maintenance, mtRNA metabolism, and translation (Rath et al., 2021). Specifically, *Hemk1* codes for an active methyltransferase that is responsible for glutamine methylation of the GGQ motif in mitochondrial release factors, which ensure efficient translation termination (Diamant et al., 2025). *Kars* encodes for an aminoacyl-tRNA synthetase, a class of enzymes that recognize and charge each essential amino acid to its respective tRNA. In mammals, the nuclear genome encodes cytosolic aminoacyl-tRNA synthetases in addition to a mostly separate set of mitochondrial aminoacyl-tRNA synthetases; however, lysyl-tRNA synthetase, encoded by *Kars*, is one of only two aminoacyl-tRNA synthetases that is involved in both nuclear and mitochondrial translation (Diodato et al., 2014; Tennakoon & Cui, 2024). In addition to being under positive selection, *Hemk1* and *Kars* are fixed or nearly fixed for *C. rutilus* ancestry in each discordant population (Table 1, Figure 5, Figure 6).

**Table 1.**
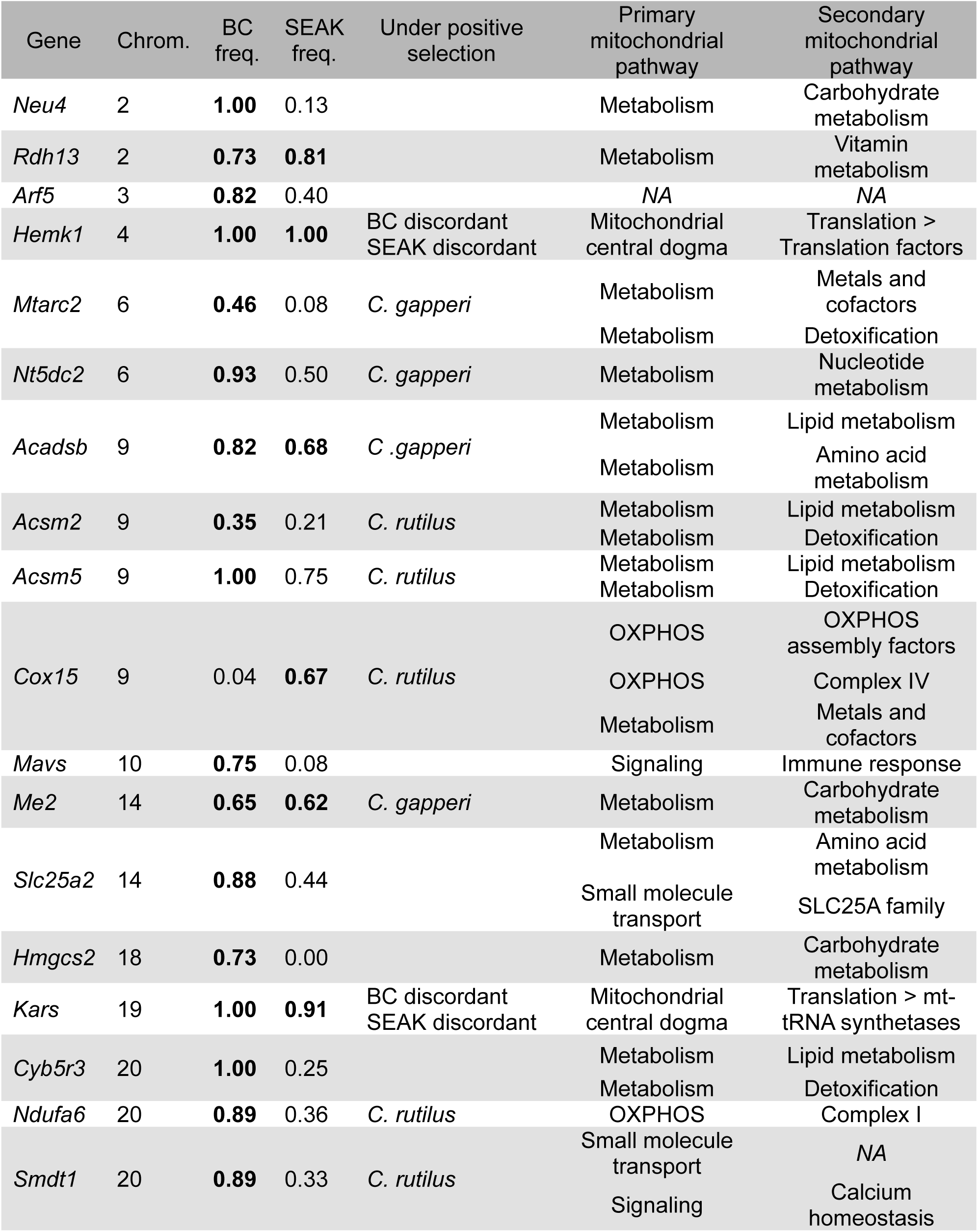
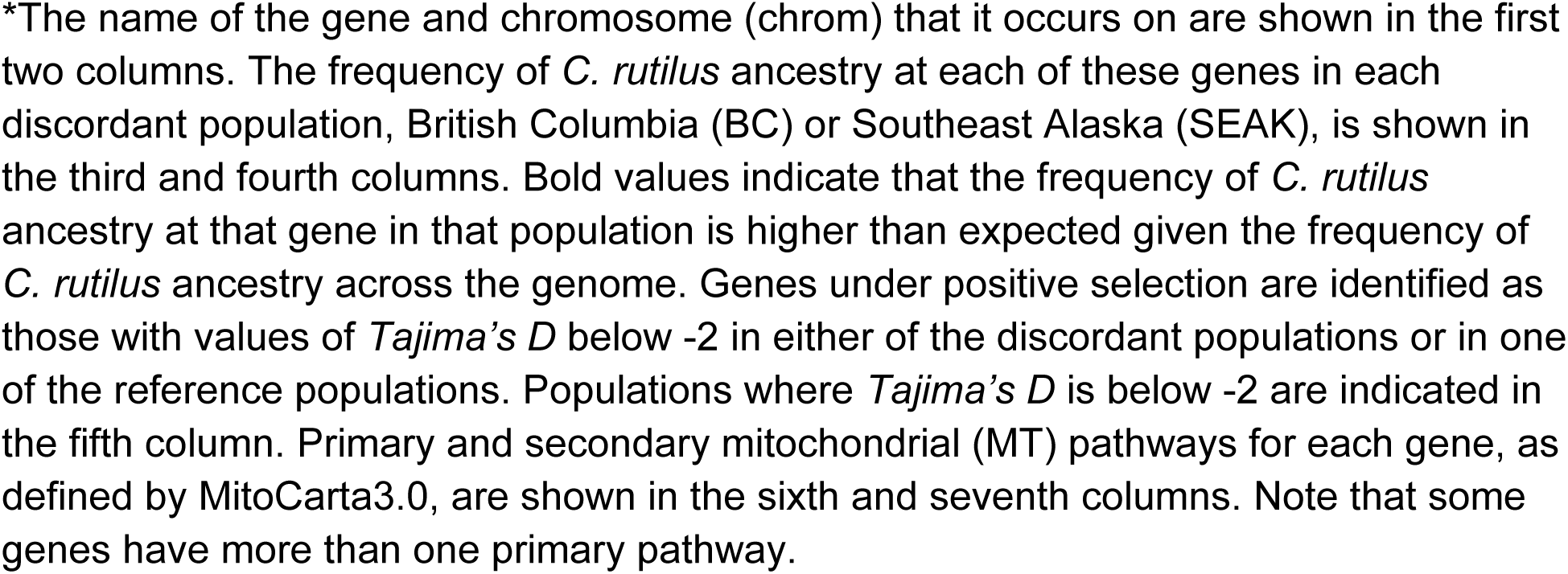
Eighteen genes identified as having significant excess *C. rutilus* ancestry in at least one of the two discordant populations.

**Figure 5.**
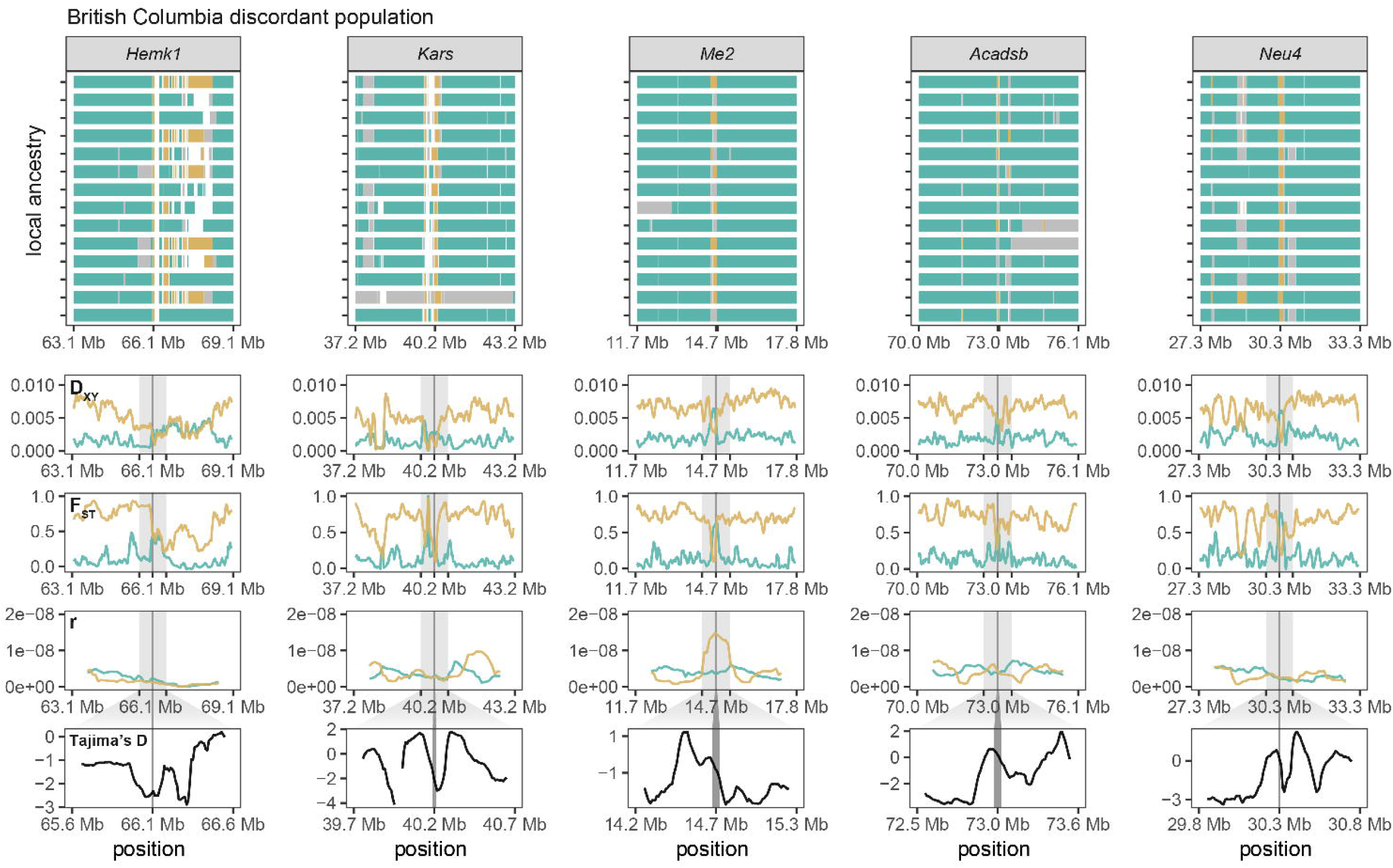
Patterns of genomic variation in the British Columbia discordant population at a subset of N-mt genes with evidence for positive selection. Each column represents a single gene and the surrounding region. The gene is centered in each panel, with its location noted by the center black tick and/or the dark grey highlighting line. The surrounding 3 Mb up- and downstream of the gene are shown for each panel, except *Tajima’s D*, which shows 500 kb up- and downstream of the gene. The range of genomic coordinates shown in the *Tajima’s D* panel are reflected by the light gray highlighting in the D_XY_, F_ST_, and recombination rate (r) panels. In the local ancestry panels (top row), each individual vole is represented by a single horizontal bar. Species ancestry at each position on the chromosome is indicated by color: *C. gapperi* = teal; *C. rutilus* = gold; heterozygous = grey. Positions with less than 95% posterior probability for any species ancestry are white. In the D_XY_ and F_ST_ panels, genetic differentiation between the British Columbia discordant population and each reference population (comparison to *C. gapperi* = teal; comparison to *C. rutilus* = gold) is summarized in 100 kb windows, slid across the region in 10 kb intervals. Recombination rate (r) is summarized for each reference population in 1 Mb windows, slid across the region in 100 kb intervals. *Tajima’s D* is summarized for the British Columbia discordant population in 100 kb windows, slid across the region in 10 kb intervals.

**Figure 6.**
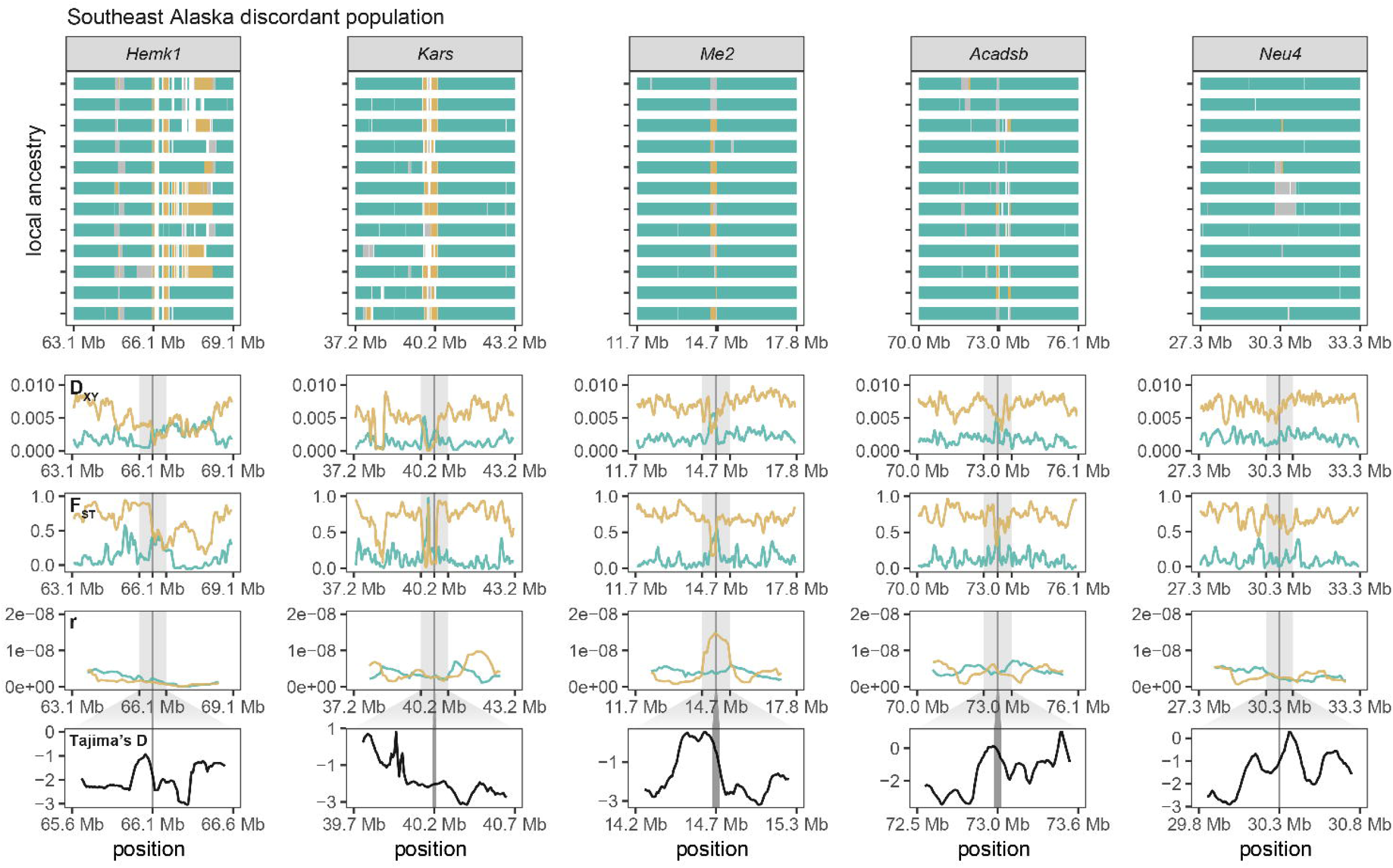
Patterns of genomic variation in the Southeast Alaska discordant population at a subset of N-mt genes with evidence for positive selection. Each column represents a single gene and the surrounding region. The gene is centered in each panel, with its location noted by the center black tick and/or the dark grey highlighting line. The surrounding 3 Mb up- and downstream of the gene are shown for each panel, except *Tajima’s D*, which shows 500 kb up- and downstream of the gene. The range of genomic coordinates shown in the *Tajima’s D* panel are reflected by the light gray highlighting in the D_XY_, F_ST_, and recombination rate (r) panels. In the local ancestry panels (top row), each individual vole is represented by a single horizontal bar. Species ancestry at each position on the chromosome is indicated by color: *C. gapperi* = teal; *C. rutilus* = gold; heterozygous = grey. Positions with less than 95% posterior probability for any species ancestry are white. In the D_XY_ and F_ST_ panels, genetic differentiation between the Southeast Alaska discordant population and each reference population (comparison to *C. gapperi* = teal; comparison to *C. rutilus* = gold) is summarized in 100 kb windows, slid across the region in 10 kb intervals. Recombination rate (r) is summarized for each reference population in 1 Mb windows, slid across the region in 100 kb intervals. *Tajima’s D* is summarized for the Southeast Alaska discordant population in 100 kb windows, slid across the region in 10 kb intervals.

## 3 | Discussion

### Strong positive selection for mitonuclear co-introgression

Incompatibilities between mitochondrial genomes and N-mt genes are expected to be one of the most common forms of DMIs, because they have different modes of inheritance yet together regulate a multitude of essential cellular processes (Burton & Barreto, 2012; Hill, 2015; Rath et al., 2021; Sloan et al., 2023). Mitonuclear incompatibilities have even been argued to be a driving force of reproductive isolation and speciation (Hill, 2015, 2019). Mitonuclear co-introgression has been put forth as a mechanism through which mitonuclear incompatibilities could be avoided in species that exhibit mitonuclear discordance, but empirical evidence for co-introgression in nature has been limited (but see Beck et al., 2015; Morales et al., 2018). Here, we use whole genome resequencing data to identify two N-mt genes, *Hemk1* and *Kars*, with a strong signal of co-introgression in two discordant populations of red-backed voles. The fixation (or near fixation) of *C. rutilus* ancestry at these genes suggests that one or more *C. rutilus* mitochondrial genes are highly incompatible with *C. gapperi* alleles at *Hemk1* and *Kars*. Further, we find evidence for positive selection on *C. rutilus* ancestry at *Hemk1* and *Kars,* suggesting recent selective sweeps in both discordant populations. These patterns are consistent with the existence of highly disadvantageous mitonuclear incompatibilities that are avoided through mitonuclear co-introgression.

While our data suggest the existence of DMIs between mitochondrial genomes and N-mt genes in this system, how the incompatibilities manifest is uncertain. Yet both of the highly co-introgressed N-mt genes are involved with mitochondrial translation, a process that relies heavily on co-adaptation of N-mt and mitochondrial genes and is predicted to be a major arena of mitonuclear incompatibility (Hill, 2015). Critically, the loading of mitochondrial tRNAs with the correct amino acids during mitochondrial translation is ensured by aminoacyl-tRNA synthetases, which are encoded in the nuclear genome (Diodato et al., 2014; Tennakoon & Cui, 2024). Incompatibilities between nuclear-encoded aminoacyl-tRNA synthetases and mitochondrial tRNAs have been identified in other systems, for example in *Drosophila* where an incompatibility between tyrosyl-tRNA synthetase and mt-tRNA^Tyr^ severely impacts oxidative phosphorylation activity and leads to delayed development and decreased female fecundity (Meiklejohn et al., 2013). In humans, reduced efficiency of mitochondrial translation is associated with a host of diseases, many of which are caused by mutations at aminoacyl-tRNA synthetases or mt-tRNAs (Richter-Dennerlein et al., 2026; Webb et al., 2020). Our identification of mitonuclear co-introgression at *Kars*, which codes for lysyl-tRNA synthetase, suggests strong selection against an incompatibility between *C. gapperi* lysyl-tRNA synthetase and *C.rutilus* mt-tRNA^Lys^ that directly impairs mitochondrial translation. The strength of observed selection against the incompatibility suggests that it directly impacts fitness through reduced viability or fertility, although further experiments will be needed to identify the exact mechanism of reduced fitness. The effect of a mitonuclear incompatibility involving *Hemk1* is less clear, as there is not a specific mitochondrial gene or RNA with which it is known to directly interact. Rather, *Hemk1* modulates the termination and fidelity of mitochondrial translation through its interaction with nuclear-encoded mitochondrial release factors (Diamant et al., 2025). The intimate role *Hemk1* plays in mitochondrial translation suggests an incompatibility could similarly impact organismal fitness through impaired energy production. In any case, while it is unclear how incompatibilities involving *Kars* and *Hemk1* manifest at the organismal level, genes involved in mitochondrial translation have been implicated in mitonuclear incompatibilities in other systems, and we add to growing a growing body of literature implicating such genes as especially susceptible to DMIs.

### Weak selection for metabolic N-mt genes

In addition to *Kars* and *Hemk1*, we identify a number of other N-mt genes with evidence for mitonuclear co-introgression, based on higher proportions of *C. rutilus* ancestry than expected given genome-wide ancestry proportions in each discordant population. These additional N-mt genes show intermediate to high frequencies of *C. rutilus* ancestry in at least one discordant population, but have not approached fixation in both (Table 1). Of the N-mt genes tested for positive selection with *Tajima’s D*, only *Kars* and *Hemk1* were significant in both discordant populations, but lack of evidence for positive selection at the other genes may be partially explained by their mixed ancestry. *Tajima’s D* is negative when the allele frequency spectrum is skewed towards many low frequency variants relative to the number of segregating sites, as is expected when a beneficial haplotype sweeps through a population (Braverman et al., 1995; Weigand & Leese, 2018). But in the case of a partial sweep of an introgressed haplotype there will essentially be two distinct haplotypes at the locus, one of each ancestry type, resulting in an excess of intermediate-frequency variants and increasing the observed value of *Tajima’s D*. In the absence of a statistic sensitive to the presence of partial sweeps of introgressed haplotypes (Harris et al., 2018; Schrider et al., 2015), we calculated *Tajima’s D* for each reference population, as a way of identifying genes under positive selection in these species. Four of these N-mt genes show evidence of positive selection in *C. gapperi* and five in *C. rutilus*, which suggests that these genes may indeed be important for survival and currently undergoing a selective sweep through one or both of the discordant populations.

Most research on mitonuclear incompatibilities has focused on DMIs that involve few genes of large effect; for example, a single N-mt gene and mitochondrial gene that, when mismatched, cause inviability or sterility (Lee et al., 2008; Meiklejohn et al., 2013; Moran et al., 2024). Such mitonuclear incompatibilities are easy to detect, both in crossing experiments and in natural settings, due to their dramatic effect on individual fitness. Our finding of multiple N-mt genes with excess, but not fixed, co-introgressed ancestry suggests that it is not uncommon for mitonuclear incompatibilities to have small effect sizes, and that these minor incompatibilities may be much more prevalent than previously appreciated. Minor mitonuclear incompatibilities could easily manifest under an environmental selective regime that favors certain genotypic combinations. Under this model, such mitonuclear incompatibilities may not be lethal, but could reduce individual fitness and be selected against over time. In the case of red-backed voles, *C. rutilus* has adapted to survive in the harsh climates of the Palearctic, where temperatures regularly drop below -40°C in the winter. In these conditions, there would be strong selection for efficient metabolism and body temperature regulation (Stevenson, 2009). If *C. rutilus* N-mt alleles provide an advantage in harsh winter conditions, especially in combination with *C. rutilus* mitochondria, there could be weak positive selection for their introgression into *C. gapperi*.

This model of environmental selection can also explain the observation of more N-mt genes with excess *C. rutilus* ancestry in British Columbia than in Southeast Alaska. Our estimates for the timing of admixture pulses in these discordant populations suggest that admixture happened in Southeast Alaska once, roughly 2,850 generations ago, while admixture in British Columbia is likely ongoing, with evidence for two pulses including one within the last 171 generations (Figure 1). This interpretation is supported by the lack of a geographic barrier separating the two species in British Columbia, and the physical isolation of the species north and south of the LeConte Bay in Southeast Alaska (Figure 2). After a single introgression event in Southeast Alaska, nuclear *C. rutilus* N-mt alleles would have been at low frequency. While mitochondrial genomes can introgress and fix quickly due to the absence of recombination and diploidy (Ballard & Whitlock, 2004; Toews & Brelsford, 2012), the discordant population in Southeast Alaska is no longer in contact with *C. rutilus*, and repeated backcrossing towards *C. gapperi* has purged most of the nuclear genome of *C. rutilus* ancestry, except at loci under strong selection. Meanwhile, continued contact in British Columbia has kept the relative proportion of *C. rutilus* ancestry in the discordant population higher, allowing more opportunity for selection to drive co-introgression of N-mt genes.

### Adaptive mitochondrial introgression despite mitonuclear incompatibilities

Despite evidence for mitonuclear incompatibilities between *C. rutilus* mitochondrial genomes and *C. gapperi* N-mt genes, there is substantial mitochondrial introgression from *C. rutilus* to *C. gapperi*, especially in British Columbia where there are fewer geographic barriers to migration than along the coastline in Southeast Alaska (Wiens & Colella, 2025). While here we only include samples from discordant red-backed voles sampled close to the center of the contact zone in British Columia, we have previously shown that mitochondrial introgression extends at least 100 km to the south (Wiens & Colella, 2025), a sizeable stretch considering that the average distance across an individual red-backed vole range is 60-145 meters (Tisell et al., 2019). It is extremely unlikely that neutral demographic- or sex-biased patterns of mating or dispersal could drive such a strong pattern of unidirectional mitochondrial introgression with such limited nuclear introgression, especially in the presence of mitonuclear incompatibilities (Bonnet et al., 2017). Further, there is no evidence that mating is biased toward *♂ C. gapperi* × ♀ *C. rutilus* crosses, as would be required under the sex-biased mating hypothesis of mitochondrial introgression (McPhee, 1977). Together, our findings point to a scenario in which *C. rutilus* mitochondria provide a high enough selective advantage that mitonuclear incompatibilities can be overcome through co-introgression of N-mt alleles.

If mitonuclear incompatibilities exist, as suggested by the presence of mitonuclear co-introgression, then individuals with intermediate admixture proportions will be rare and successful mitochondrial introgression events uncommon, both of which are supported by the patterns of genetic variation observed across the landscape (Hill et al., 2019; Sloan et al., 2017). In British Columbia, where there are no geographic barriers to contact, nuclear ancestry turns over rapidly from full *C. rutilus* to the discordant population, where individuals have between 6.4% to 9.3% *C. rutilus* nuclear ancestry. This narrow range of low ancestry proportions suggests repeated backcrossing towards *C. gapperi*, with intermittent input from *C. rutilus*. The absence of early stage hybrids with intermediate admixture proportions (e.g., F1s or F2s), even near the center of the contact zone, suggests that interspecies matings are rare and/or intermediate admixture proportions result in unfit genotypes, as would be expected when postzygotic reproductive barriers exist in the form of DMIs (Hill et al., 2019; Maheshwari & Barbash, 2011; Thompson et al., 2023). The mitochondrial phylogeny further supports a scenario of continued, intermittent pulses of ancestry from *C. rutilus*. Introgressed *C. rutilus* mitochondrial haplotypes in British Columbia form two distinct clades, each with essentially no variability, indicating at least two independent introgression events in the very recent past (Figure 2). In Southeast Alaska, on the other hand, introgressed *C. rutilus* mitochondrial haplotypes display variation but share a common ancestor, supporting a single, distant introgression event, occurring before the species were isolated from each other across the LeConte Bay. While all introgressed mitochondrial haplotypes in British Columbia and Southeast Alaska form a single, well-supported clade (96% bootstrap support), there is only one position in the mitogenome (16S rRNA position 859) that supports this node, shared by all introgressed *C. rutilus* mitogenomes to the exclusion of non-introgressed *C. rutilus* mitogenomes. Interestingly, all non-introgressed *C. rutilus* mitochondrial haplotypes have a T at this position, while introgressed *C. rutilus* haplotypes have a C at this position, which is shared by *C. gapperi* mitochondrial haplotypes. The functional significance of a T→C mutation at this position is unclear, but the fact that all introgressed mitochondrial haplotypes share this substitution with *C. gapperi* suggests that it may be important for successful introgression of *C. rutilus* mitochondria against *C. gapperi* nuclear backgrounds.

For introgression of *C. rutilus* mitochondria to repeatedly occur in the face of mitonuclear incompatibilities, they must provide a strong selective advantage. Given that *C. rutilus* has persisted in the high Arctic for the last two million years, one possible explanation is that *C. rutilus* mitochondria are involved in adaptations that confer increased survival through the winter. Red-backed voles do not hibernate or go into torpor in the winter, unlike some other high-latitude mammals; instead, they remain metabolically active in the subnivium, where they are able to forage for resources through the winter (Jolly et al., 2022; Stevenson, 2009). Previous studies of *C. rutilus* have found various physiological adaptations to extreme winter conditions, including an increase in maximum metabolic rate (Rosenmann et al., 1975) and an overall reduction in body mass across tissue types, with the exception of brown adipose tissue (BAT), which increases in relative mass in the winter (Zuercher et al., 1999). BAT is the center for non-shivering thermogenesis in mammals, a process that is driven by the loss of energy as heat during mitochondrial oxidative phosphorylation. Specifically, upregulation of *Ucp1* in BAT uncouples ATP synthesis from oxidative phosphorylation and instead allows the proton gradient to dissipate as heat (Chouchani et al., 2019; Steensels & Ersoy, 2019). It is possible that *C. rutilus* mitochondria have co-evolved with N-mt genes to increase heat production through non-shivering thermogenesis, which could be facilitated by changes in gene expression, metabolic efficiency, or mitochondrial density in BAT, and which would increase the likelihood of winter survival (Aquilano et al., 2023; Chouchani et al., 2019). If *C. rutilus* physiological adaptations to extreme cold are facilitated by mitochondrial evolution, mitonuclear co-introgression may well be adaptive for *C. gapperi* populations that have expanded north into the same environment. In line with this idea, three of the N-mt genes with evidence for co-introgression (*Acadsb*, *Acsm2*, and *Acsm5*) are involved in β-oxidation, a process by which fatty acids are broken down to produce energy released as heat during non-shivering thermogenesis (Houten et al., 2016; Steensels & Ersoy, 2019). Notably, every individual in both discordant populations is either heterozygous or homozygous for *C. rutilus* ancestry at *Acadsb*, a gene that has previously been found to be upregulated along with *Ucp1* in the BAT of mice exposed to prolonged cold (Amor et al., 2024). While *Ucp1* (also known as *Slc25a7*) doesn’t show evidence of co-introgression here, there is evidence of co-introgression for a related gene, *Slc25a2*, which like *Ucp1* belongs to the *Slc25* mitochondrial carrier gene family (Jones et al., 2024). Relatively little is known about *Slc25a2*, but its relation to *Ucp1* hints at a possible role in non-shivering thermogenesis. Overall, co-introgression of mitochondria and multiple N-mt genes associated with non-shivering thermogenesis suggests introgression is beneficial for northern *C. gapperi* populations. In fact, adaptive introgression of *C. rutilus* mitochondria along thermal regimes has been previously proposed in an independent contact zone in Europe, where *C. rutilus* mitochondria have introgressed into populations of *C. glareolus* (Boratyński et al., 2011, 2014, 2016; Marková et al., 2020; Šíchová et al., 2014). While our data cannot confirm non-shivering thermogenesis as the mechanism mediating adaptive mitochondrial introgression, what is known about *C. rutilus* winter physiology suggests that selection for winter survival has shaped the evolution and introgression of mitochondrial genomes in this system.

## Conclusions

Mitonuclear co-introgression has been hypothesized as a mechanism that could facilitate mitochondrial introgression, but there are exceedingly few examples of this phenomenon (Beck et al., 2015; Morales et al., 2018; Sloan et al., 2017). One explanation for the paucity of evidence for mitonuclear co-introgression is that population-level whole genome resequencing data is needed to identify deviations in ancestry proportions across thousands of N-mt genes, which has not been feasible for most systems until recently. Here, we show that selection has driven *C. rutilus* mitochondrial introgression into northern *C. gapperi* populations and that mitonuclear incompatibilities have been overcome through co-introgression at N-mt genes. Our findings support the hypothesis that mitonuclear incompatibilities between species are common, given the intimate relationship between mitochondrial genomes and N-mt genes, but can be circumvented through mitonuclear co-introgression. While genomic data is useful for detecting selection at the molecular level, future work should incorporate experimental physiology to test hypotheses about the adaptive benefit of introgressed mitonuclear networks. Overall, our findings highlight the importance of considering the interplay between nuclear and mitochondrial genomes when investigating species relationships and suggest that mitonuclear interactions can play an important role in shaping outcomes of hybridization and speciation.

## 4 | Materials and Methods

### Library preparation and whole genome resequencing

Frozen tissue (liver or kidney) subsamples were loaned from three natural history collections: University of Kansas Biodiversity Institute (KU), University of New Mexico Museum of Southwestern Biology (MSB), and University of Alaska Museum of the North (UAM). See Supplementary File 1 for full specimen metadata. The sequenced reference populations of each species (*C. rutilus* and *C. gapperi*) consisted of 21 and 13 specimens, respectively. All reference individuals were collected within a single two-day span and from a single locality, both of which occur outside the known contact zone in British Columbia (Wiens & Colella, 2025). Our sampling also included 14 discordant voles from British Columbia, 12 discordant voles from Southeast Alaska, 4 non-reference *C. rutilus* from British Columbia, 6 non-reference *C. rutilus* from Southeast Alaska, and 2 non-reference *C. gapperi* from Southeast Alaska. In total, DNA was extracted from frozen tissue subsamples of 72 *Clethrionomys* specimens and one *Alticola lemminus* specimen, as an outgroup, following a modified magnetic bead-based extraction protocol (Rohland & Reich, 2012). Libraries were fragmented enzymatically to an average insert size of 434 bp and prepared for whole genome resequencing with the xGen™ DNA Lib Prep EZ kit from IDT. Libraries were sequenced either on an Illumina NovaSeq X Plus at the Oklahoma Medical Research Foundation Clinical Genomics Center or an Illumina NextSeq2000 at the University of Kansas Genome Sequencing Core, in all cases generating 150 bp paired-end reads. On average, 188.5 million paired-end reads were generated per individual.

### Bioinformatic processing

Raw sequencing reads were trimmed and filtered for quality using *fastp* v0.23.4 (Chen, 2023). Read ends were trimmed using 4 bp sliding windows until the average phred score was >=20. Trimmed reads with >10% of the bases below a phred score of 20 or with >5 uncalled bases were discarded. Passing reads whose mate did not also pass quality filters were discarded, such that only paired reads were retained. Filtered reads were aligned to the *C. rutilus* reference genome (GCA_040207285.2) using the *mem* algorithm in *bwa* v0.7.18 (Li & Durbin, 2009). On average, individual read depth was 20.1X (see Supplementary File 1 for read depth per individual).

### Mitogenome phylogenies

We extracted full mitogenome sequences from individual alignments using the *consensus* command in *samtools* v1.21 (Danecek et al., 2021). Initial inspection revealed large stretches of missing data for individuals with *C. gapperi* mitogenomes, so we realigned those individuals to a *C. gapperi* mitogenome reference (GenBank: [NC_068811](https://www.ncbi.nlm.nih.gov/nuccore/2386966345)) and re-extracted consensus mitogenomes. Average mitogenome depth was 18,995X for *C. rutilus* and 20,689X for *C. gapperi*. We then re-aligned all consensus mitogenome sequences to each other. We built a maximum-likelihood mitogenome phylogeny using *iqtree* v2.3.6 (Minh et al., 2020), implementing model testing to find the best model of sequence evolution (Kalyaanamoorthy et al., 2017), with 1,000 ultrafast bootstraps.

### Patterns of hybrid ancestry in the nuclear genome

To provide an initial characterization of ancestry patterns in the nuclear genome, we generated a dataset of reduced linkage, high-quality SNPs. We called biallelic SNPs for all 73 individuals with *bcftools* v1.21 (Danecek et al., 2021), filtering for minimum mapping quality and base quality of 30, removing indels, and keeping only variant sites. We filtered for quality by removing SNPs with extreme (defined as above or below two standard deviations) read position bias (RPBZ), mapping quality bias (MQBZ), mapping quality vs strand bias (MQSBZ), base quality bias (BQBZ), or soft-clip length bias (SCBZ). Sites with abnormally low or high read depth may result in erroneous genotype calls, so we filtered SNPs for minimum and maximum depth. To account for slight variation in total reads per individual, we calculated the 95th quantile of depth for each individual and set sites above that value to missing data. We set a minimum read depth of five for all individuals. We then filtered for SNPs with non-missing data in at least 2 individuals and pruned SNPs for a minimum distance of 10 kb, resulting in 234,874 SNPs with 15.9% overall missing data. From those SNPs, we used *triangulaR* v0.0.1 to identify 25,375 AIMs, defined as SNPs with an allele frequency difference of 1 (i.e., fixed differences) between the reference *C. rutilus* and *C. gapperi* populations (Wiens et al., 2025). We used those AIMs to estimate hybrid ancestry for each individual by calculating hybrid index and interclass heterozygosity and visualizing results on a triangle plot.

### Demographic history

We estimated demographic histories for each species using the sequentially Markov coalescent model implemented in *smc++* v1.15.5 (Terhorst et al., 2017). We called biallelic SNPs for each reference population separately using *bcftools*, filtering for minimum mapping quality and base quality of 30, removing indels, and keeping only variant sites. In total, 56.1 Mb were variant in *C. rutilus* and 80.7 Mb were variant in *C. gapperi*. *smc++* takes as input SNPs for a single reference population and assumes all other sites are invariant. To account for this, we implemented a number of quality filters for each reference population by converting non-passing sites to missing data to avoid such sites being treated as invariant. To filter for genotype quality, sites with extreme (defined as above or below two standard deviations) values of RPBZ, MQBZ, MQSBZ, BQBZ, or SCBZ were set to all missing data. Sites with read depth below 10 or above the 95th quantile of depth within each individual were set to missing data. We then converted sites with missing data for two or more individuals to all missing data. It is difficult to detect SNPs in regions of the genome with low read depth (such as centromeres), so to avoid treating these regions as long runs of homozygosity, we created a mask of positions to be marked as missing data, as per the recommendation of the *smc++* manual. To create the mask, we used *pandepth* v2.26 to calculate average depth in 100 bp sliding windows, and masked windows with average depth <2 across all individuals in a reference population (Yu et al., 2024). For each species, we prepared SNP data for *smc++* using the command *vcf2smc* and randomly chose 10 individuals as the “distinguished lineages”. We then ran *estimate* to fit cubic spline models of effective population size using the composite likelihood of the runs based on each distinguished lineage. We bounded demographic history inference between the last 10 and 1,000,000 generations, specified ancestral alleles as unknown, assumed a mutation rate of 8.7 x 10^-9^ (Wang et al., 2023), and increased the *ftol* parameter to 1e-3 to prevent model overfitting. For visualization, we converted generations to years by assuming an average of 1.5 generations per year (Banfield, 1974).

### Recombination rates

We estimated fine-scale recombination rates for each species using *pyrho* v0.2.0, a linkage disequilibrium (LD) based method which uses demography-informed composite-likelihood to estimate per-generation recombination rates based on population genomic data (Kamm et al., 2016; Spence & Song, 2019). Using the same SNP data as for demographic history estimation, we first computed a lookup table with the *make_table* command, providing the demographic history inferred from *smc++* and assuming the same mutation rate. After generating lookup tables, we ran the *hyperparam* command to optimize the main hyperparameters of *pyrho* (e.g., window size and block penalty) for our data. We then estimated recombination rates by running the *optimize* command with the preferred hyperparameters. To test whether the difference in sample size for the *C. rutilus* (N = 21) and *C. gapperi* (N = 13) reference populations impacted results, we re-estimated *C. rutilus* recombination rates with a random downsample of 13 individuals. To examine variation in recombination rate across the genome, we calculated mean recombination rates in 1 Mb sliding windows with a 100 kb step-size using *bedtools* v2.31.1 (Quinlan & Hall, 2010). For *C. rutilus*, the estimated recombination rates in sliding windows using 21 samples and 13 samples were highly correlated (Spearman’s ρ = 0.78, *P* < 2.2 × 10^-16^), so we used the recombination rates estimated with the full *C. rutilus* dataset (N = 21) moving forward.

### Genome-wide summary statistics

We used *pixy* v2.0.0 (Korunes & Samuk, 2021) to estimate genetic variation across the genome for each population using a sliding window approach for multiple summary statistics (π, F_ST_, D_XY_, *Tajima’s D*). To avoid biased estimates, we included variant and invariant sites for each population (Korunes & Samuk, 2021). We called genotypes for all sites on the 27 autosomes using *bcftools*. While calling genotypes we filtered for a minimum base quality of 30, applied the HWE assumption only within populations, and skipped indels. We then removed sites with more than two alleles and removed sites with extreme (defined as above or below two standard deviations) values of RPBZ, MQBZ, MQSBZ, BQBZ, or SCBZ. Genotypes with read depth below five and above the 95th quantile of the depth distribution within each individual were set to missing. We then removed sites with non-missing genotypes for one or fewer individuals, resulting in 2.05 Gb of total called sites (1.98 Gb invariant, 65.6 Mb variant) with 12.7% missing data on average per individual. We calculated summary statistics across the genome for each population in *pixy* using two sets of sliding windows: 1 Mb windows with a 100 kb step-size and 100 kb windows with a 10 kb step-size.

### Local ancestry inference and the timing of secondary contact

We inferred local ancestry across the nuclear genomes of the two discordant populations (BC and SEAK) using *Ancestry_HMM*, which implements a hidden Markov model and uses read counts and ancestry tract lengths to simultaneously estimate local ancestry and time since admixture (Medina et al., 2018). Since *Ancestry_HMM* was optimized for read counts instead of called genotypes and does not require invariant sites, we used a modified version of the same dataset used for *pixy*. Specifically, we implemented the same quality filters up to the point where we filtered for depth. At this point, we used *bcftools* to filter for variable, biallelic SNPs and removed sites that were above the 95th quantile for total read depth across all samples, as such sites may represent paralogous loci or have high repeat content. To infer local ancestry in admixed populations, *Ancestry_HMM* relies on reference panels of the groups that supplied the ancestry to identify ancestry informative markers. So, we further filtered our sites for <25% missing data within each reference population (i.e., a site had to be non-missing in at least 16 of 21 reference *C. rutilus* and 10 of 13 reference *C. gapperi*). We then prepared our dataset for local ancestry inference using a slightly modified (to account for allelic depth being in the 4th column of the genotype line of the vcf) version of the python script (*vcf2ahmm.py*) provided by the authors of *Ancestry_HMM*. Discordant individuals had majority *C. gapperi* nuclear ancestry, so we specified an average recombination rate of 3.34664e-09 (as estimated for *C. gapperi*). We filtered for sites with a 10% minimum allele frequency difference between the reference populations, and thinned for a minimum distance of 1 kb between sites.

For each discordant population, we built and assessed the fit of two models: a model consisting of a single pulse of *C. rutilus* ancestry into the discordant population (one-pulse model) and a model consisting of two, distinct pulses, both from *C. rutilus* (two-pulse model). For each model, we provided initial estimates of global ancestry proportions as inferred with *triangulaR*, specified that pulses must occur within the last one million generations, and provided the estimated effective population size of *C. gapperi* (N_e_ = 90,166). We provided an initial estimate of contact at 100 generations ago for the one-pulse model and added a less recent pulse at 1,000 generations ago for the two-pulse model, allowing the timing of the pulses to be optimized during fitting of the admixture model. We also estimated the relative proportion of ancestry contributed by each *C. rutilus* pulse in the two-pulse model. We performed 1,000 bootstraps with a block size 1,000 for the one-pulse models and 50 bootstraps (due to computational restraints) with the same block size for the two-pulse models.

### Ancestry analysis at nuclear-encoded mitochondrial genes

Leveraging local ancestry posterior probabilities as inferred by *Ancestry_HMM*, we called ancestry states (i.e., homozygous *C. rutilus*, heterozygous, or homozygous *C. gapperi*) across the nuclear genome of each discordant individual. Sites with less than 95% posterior probability support for any ancestry state were left unclassified. To test for excess *C. rutilus* ancestry at N-mt genes, we binned regions of the nuclear genome with the following approach. First, we identified N-mt genes with annotations in the *C. rutilus* (GCA_040207285.2) reference genome, as classified by Mouse MitoCarta3.0 (Rath et al., 2021). Next, we used the recombination maps generated for *C. gapperi* to calculate the length of each N-mt gene, finding an average length of 0.0086 cM. We then divided the nuclear genome into 0.0086 cM windows and removed any window that overlapped with an N-mt gene, resulting in a set of genomic windows excluding N-mt genes. For each discordant population, we calculated the average proportion of *C. rutilus* ancestry across individuals in each N-mt gene and genomic window. For the genomic windows, we used linear regression to test for an association between recombination rate and the proportion of *C. rutilus* ancestry.

We used two approaches to test the BC and SEAK discordant populations for biased introgression of *C. rutilus* ancestry at N-mt genes. In the first approach, we tested whether N-mt genes were enriched for *C. rutilus* ancestry as compared to the genomic windows using the Kolmogorov-Smirnov test as implemented in R. In the second approach, we took 10,000 random samples of 1,124 genomic windows (the same as the number of N-mt genes) and calculated the average *C. rutilus* ancestry in each sample. This approach provided a null distribution of the average *C. rutilus* ancestry expected in a random sample of 1,124 genomic windows for each discordant population. We then calculated the 2.5% and 97.5% quantiles of those distributions, against which we tested the observed proportion of *C. rutilus* ancestry at N-mt genes.

### Tests of selection for heterospecific ancestry

After finding evidence for biased introgression of *C. rutilus* ancestry at N-mt genes as a whole, we further tested each N-mt gene independently for excess *C. rutilus* ancestry as compared to the genomic windows. We used a slightly modified version of the CAnD test described by McHugh et al. (2016), which facilitates testing for differences in mean values among correlated groups. Specifically, for each N-mt gene, we tested the distribution of *C. rutilus* ancestry proportions across individuals against the distribution of mean *C. rutilus* ancestry in the rest of the genome. We applied a Bonferroni correction to account for multiple hypothesis testing across the 1,124 N-mt genes.

After identifying a subset of N-mt genes that showed evidence for biased introgression in either or both of the discordant populations, we tested for evidence of selective sweeps of *C. rutilus* ancestry at those genes. To do so, we calculated *Tajima’s D* for each gene in each discordant population using *pixy* and the same genomic dataset as for the sliding window-based estimates of *Tajima’s D*, described above. We considered values of *Tajima’s D* below -2 as significant for positive selection for *C. rutilus* ancestry in a discordant population. This approach can detect complete or near complete selective sweeps, but will fall short at detecting positive selection for incomplete sweeps. Therefore, we also tested these genes for evidence of positive selection within each reference population, which provides some information about which genes are under selection in the two species, despite limitations for the discordant populations.

## Data and resource availability

All specimens used in this study are reported in the Specimen Appendix (Supplementary File 1). All sequencing raw reads will be uploaded to the NCBI Sequence Read Archive (SRA) pending manuscript acceptance. All code used for data analysis is available at https://github.com/omys-omics/RBV_WGS.

## Acknowledgements

We thank the American Philosophical Association for a Lewis and Clark grant that supported field work, the American Genetic Association for an EECG Research Award in support of whole genome resequencing, and the KU Center for Genomics for a Seed Grant that further supported sequencing efforts. Further support for this project was provided by the NSF (DBI-2100955). We thank multiple field crews who assisted in collecting red-backed vole museum specimens in Southeast Alaska and British Columbia, including Bailey Dixon, Alex Hey, Errol Hooper, Danny Ibañez, Dianna Krejsa, Danielle Land, and Robert Nofchissey. We thank the British Columbia Ministry of Water, Land and Resource Stewardship and the Alaska Department of Fish & Game for scientific collecting permits. We thank the curators and collection managers at the University of New Mexico Museum of Southwestern Biology (Jon Dunnum, Mariel Campbell, Joe Cook), University of Alaska Museum of the North (Kyndall Hildebrandt, Mallory Gulbranson, Aren Gunderson, Link Olsen) and University of Kansas Biodiversity Institute (Dianna Krejsa) for tissue loans of red-backed vole specimens. The availability of museum red-backed specimens, especially from the reference populations of each species, was made possible by the field and museum archival efforts of the Center for Integrative Inventories of Biomes of the Arctic (CIIBA, NSF DEB-1258010) and the Beringian Co-Evolution Project (NSF DEB-0196095). We thank the Oklahoma Medical Research Foundation Clinical Genomics Center and University of Kansas Genome Sequencing Core for Illumina sequencing. Data analysis was performed on the HPC facilities operated by the Center for Research Computing at the University of Kansas supported in part through the National Science Foundation MRI Award #2117449. We thank the Canadian BioGenome Project for generating and making available a *Clethrionomys rutilus* chromosome-level reference genome.

## Author contributions

BJW and JPC conceptualized and funded the study and led field work to collect red-backed vole specimens. BJW extracted DNA and prepared whole genome resequencing libraries, performed data analysis, and wrote the first version of the manuscript. JPC provided conceptual and editorial input throughout.

