## Supplementary File 2 for "Mitochondrial introgression in North American red-backed voles is facilitated by co-introgression at nuclear-encoded mitochondrial genes"

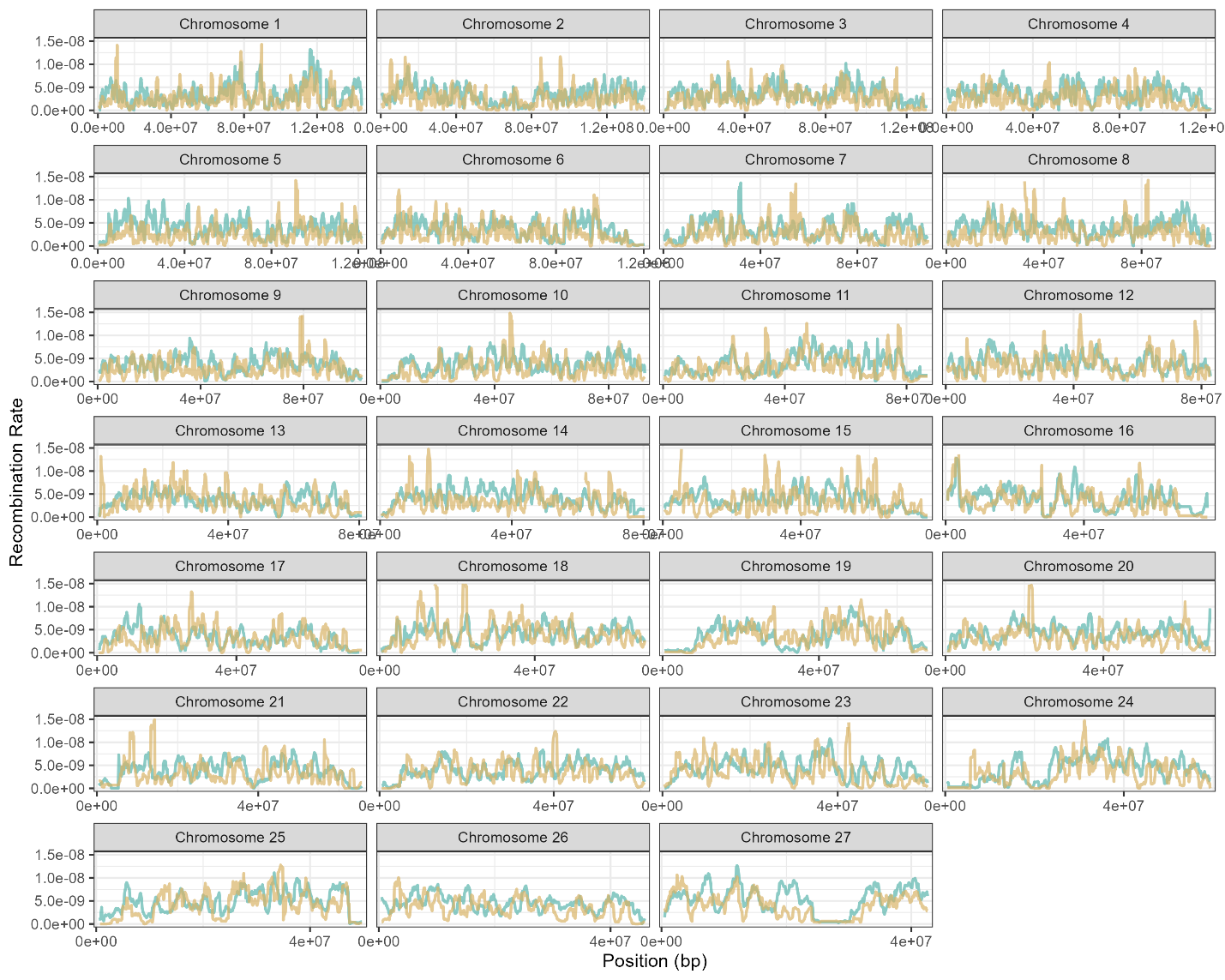

**Figure S1.** Estimated recombination rates (r) along each autosome for each species (*C. gapperi* = teal; *C. rutilus* = gold). Recombination rate is summarized in 1 Mb windows, slid across each chromosome in 100kb intervals.

**Figures S2-S19. The following pages show patterns of genomic variation in the British Columbia discordant population at each N-mt gene from Table 1.** Each column represents a single gene and the surrounding region. From top to bottom, the panels show local ancestry, D_XY_, F_ST_, recombination rate, and *Tajima’s D*. The gene is centered in each panel, with its location noted by the center black tick and/or the dark grey highlighting line. The surrounding 3 Mb up- and downstream of the gene are shown for each panel, except for *Tajima’s D*, which shows the surrounding 500 kb up- and downstream of the gene. The range of genomic coordinates shown in the *Tajima’s D* panel are reflected by the light gray highlighting in the D_XY_, F_ST_, and recombination rate (r) panels. In the local ancestry panels, each individual vole is represented by a single horizontal bar. Species ancestry at each position of the chromosome is indicated by color: *C. gapperi* = teal; *C. rutilus* = gold; heterozygous = grey. Positions with less than 95% posterior probability for any species ancestry are white. In the D_XY_ and F_ST_ panels, genetic differentiation between the British Columbia discordant population and each reference population (comparison to *C. gapperi* = teal; comparison to *C. rutilus* = gold) is summarized in 100 kb windows, slid across the region in 10 kb intervals. Recombination rate (r) is summarized for each reference population in 1 Mb windows, slid across the region in 100 kb intervals. *Tajima’s D* is summarized for the British Columbia discordant population in 100 kb windows, slid across the region in 10 kb intervals.

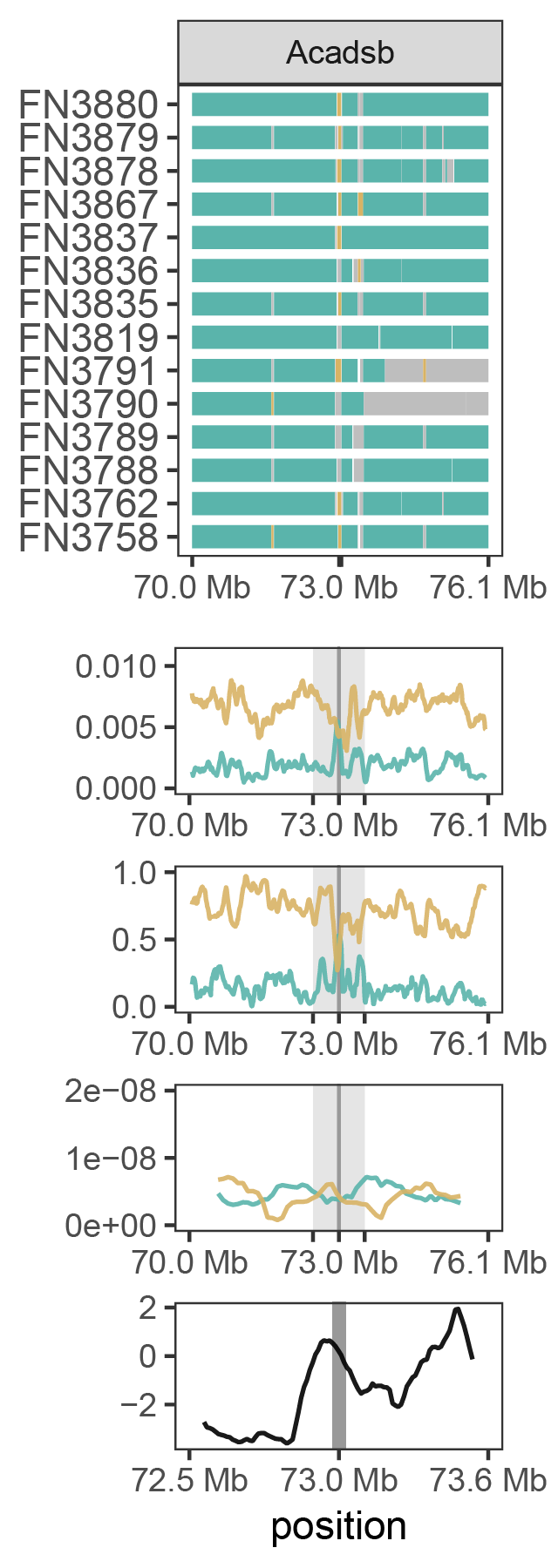

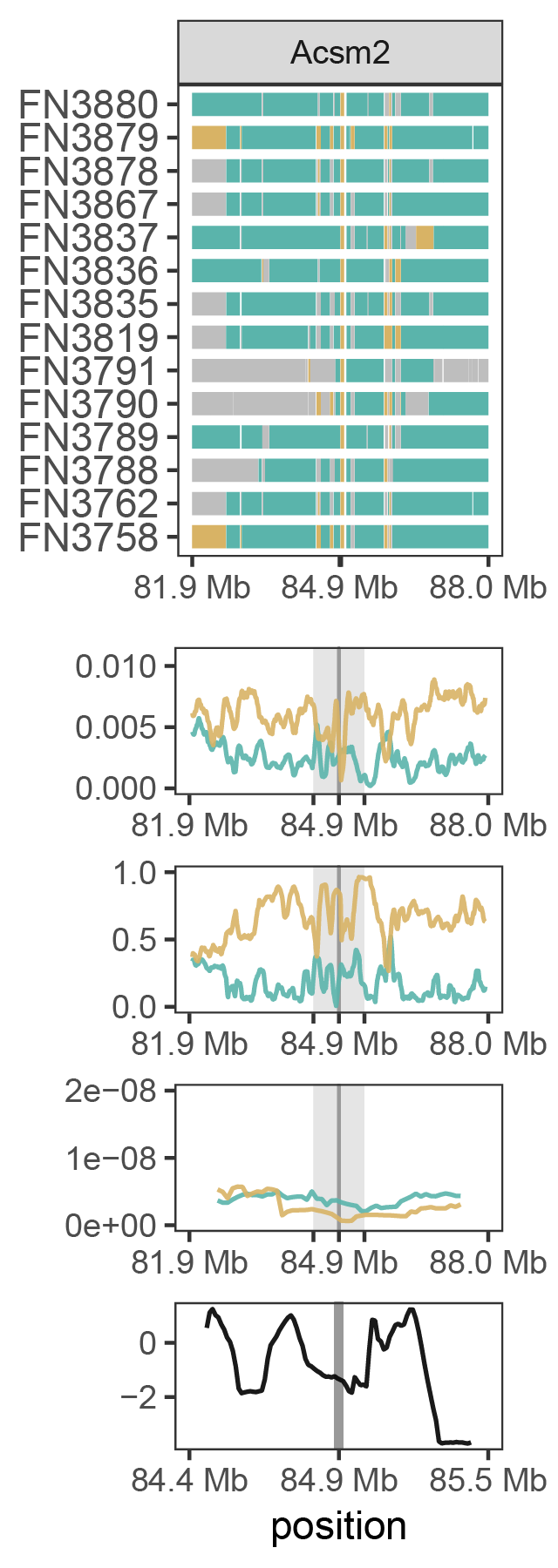

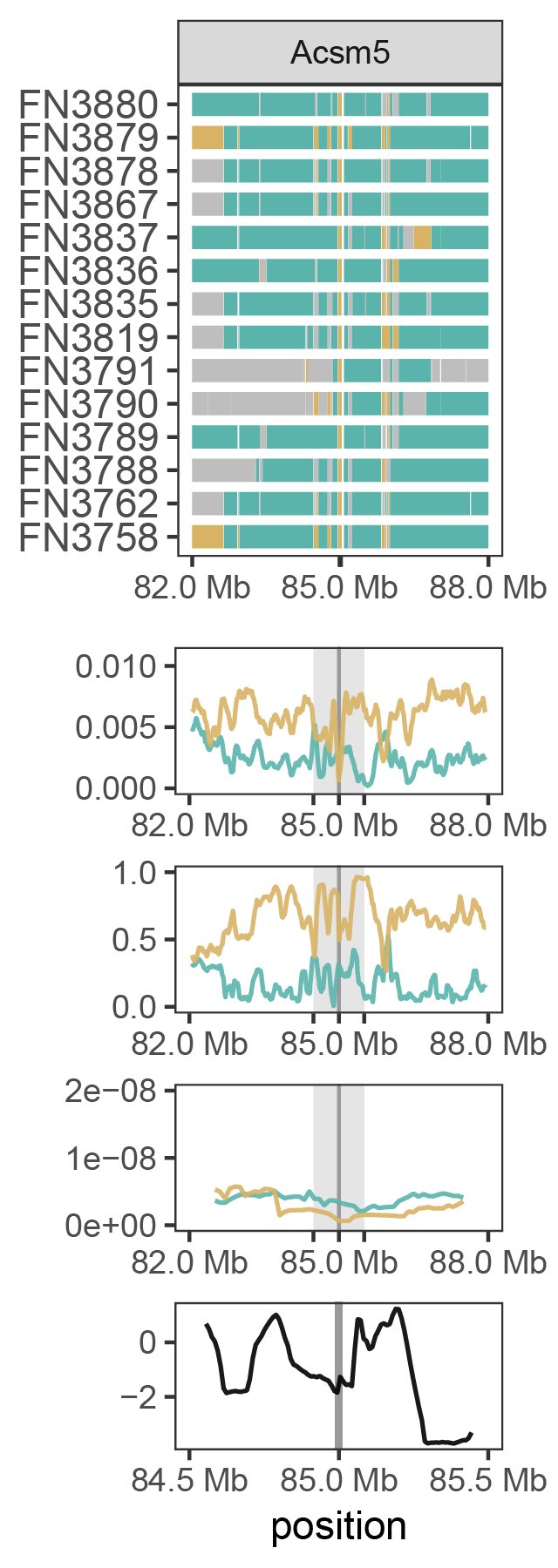

**Figures S2-S4.**

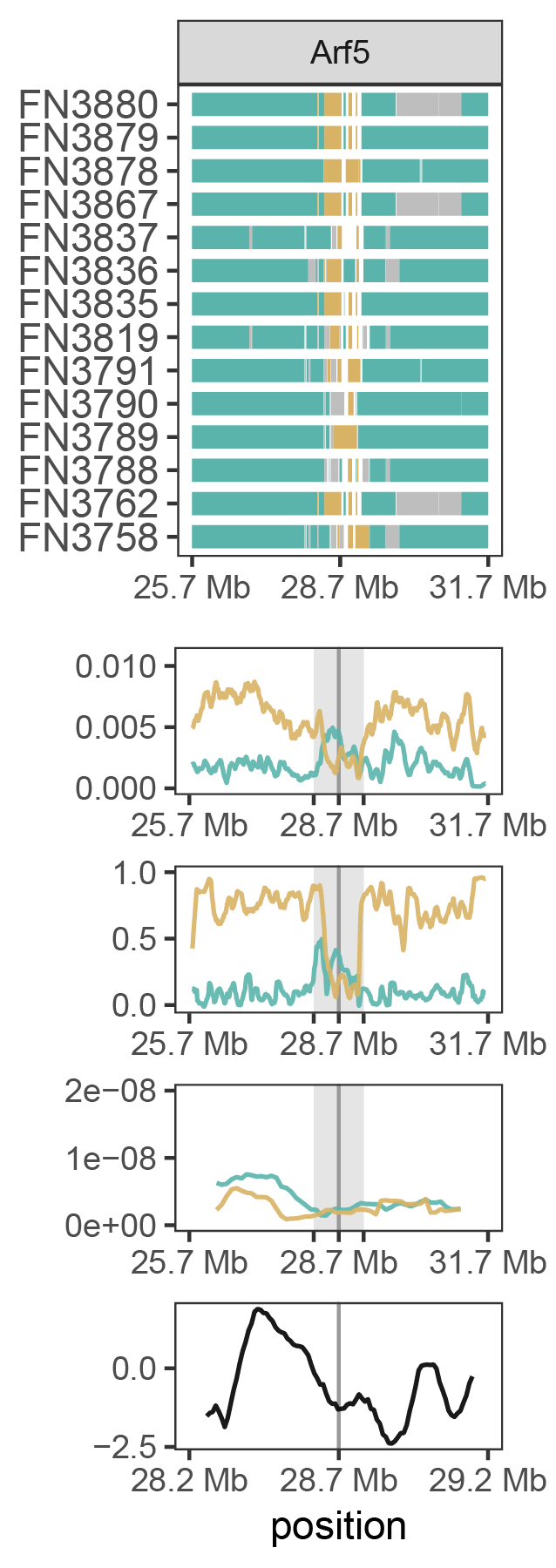

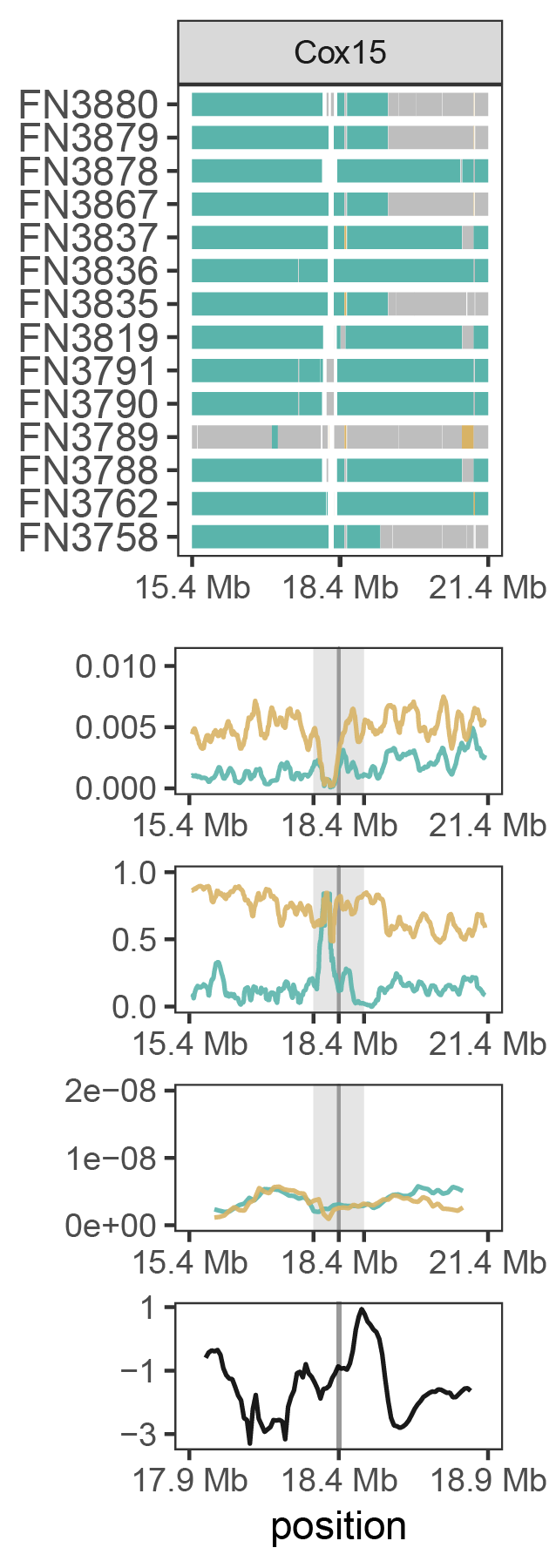

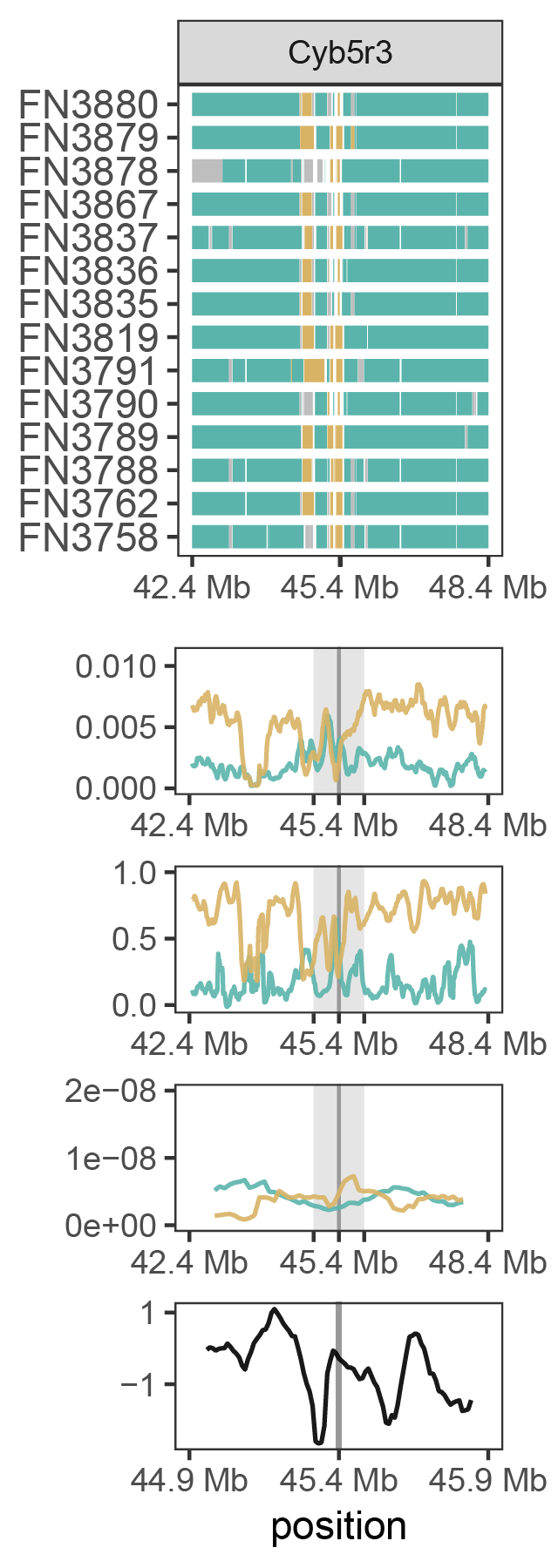

**Figures S5-S7.**

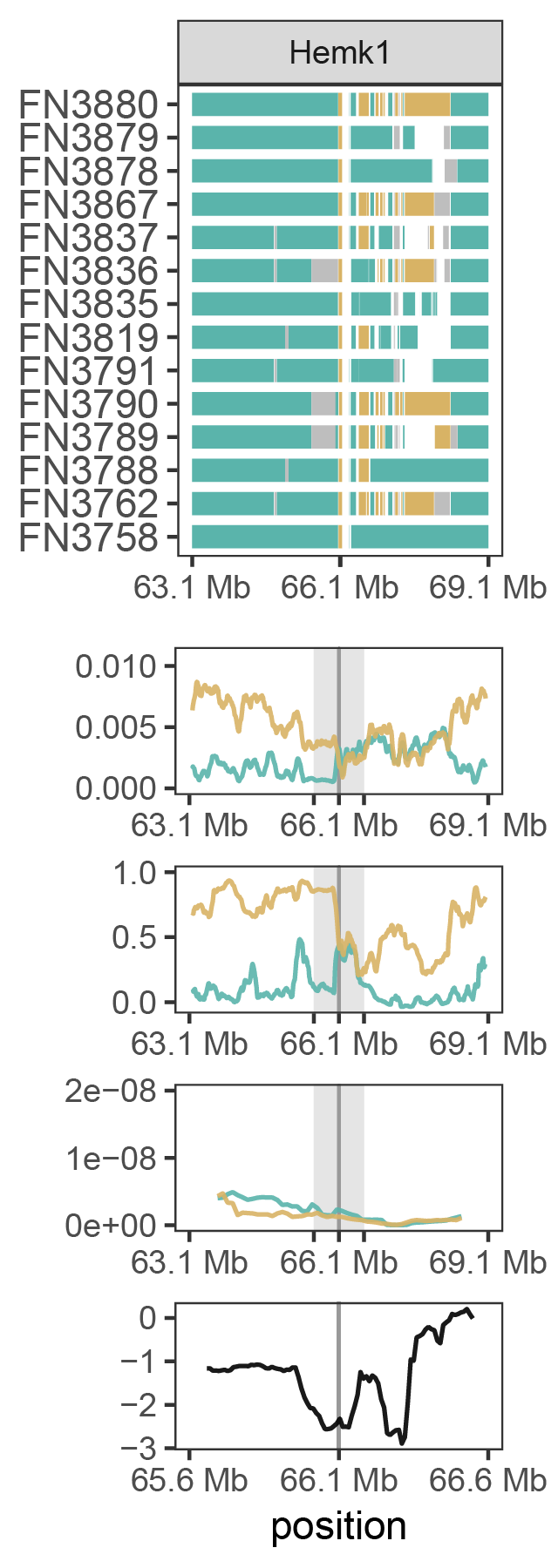

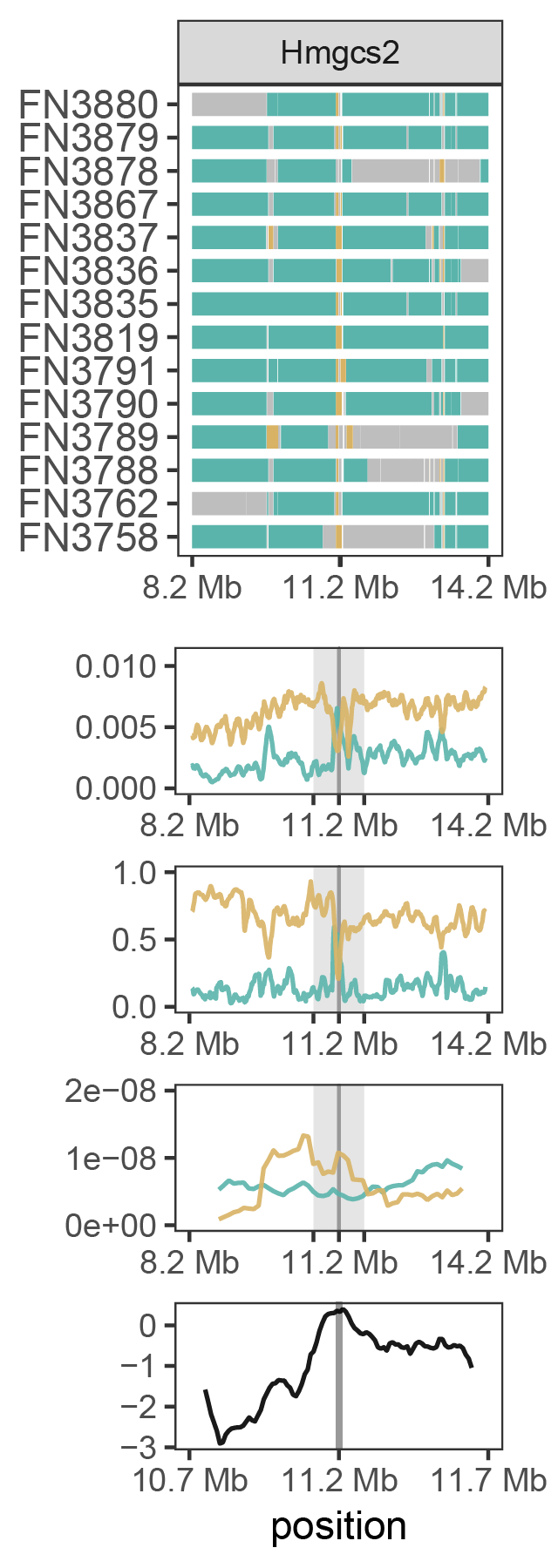

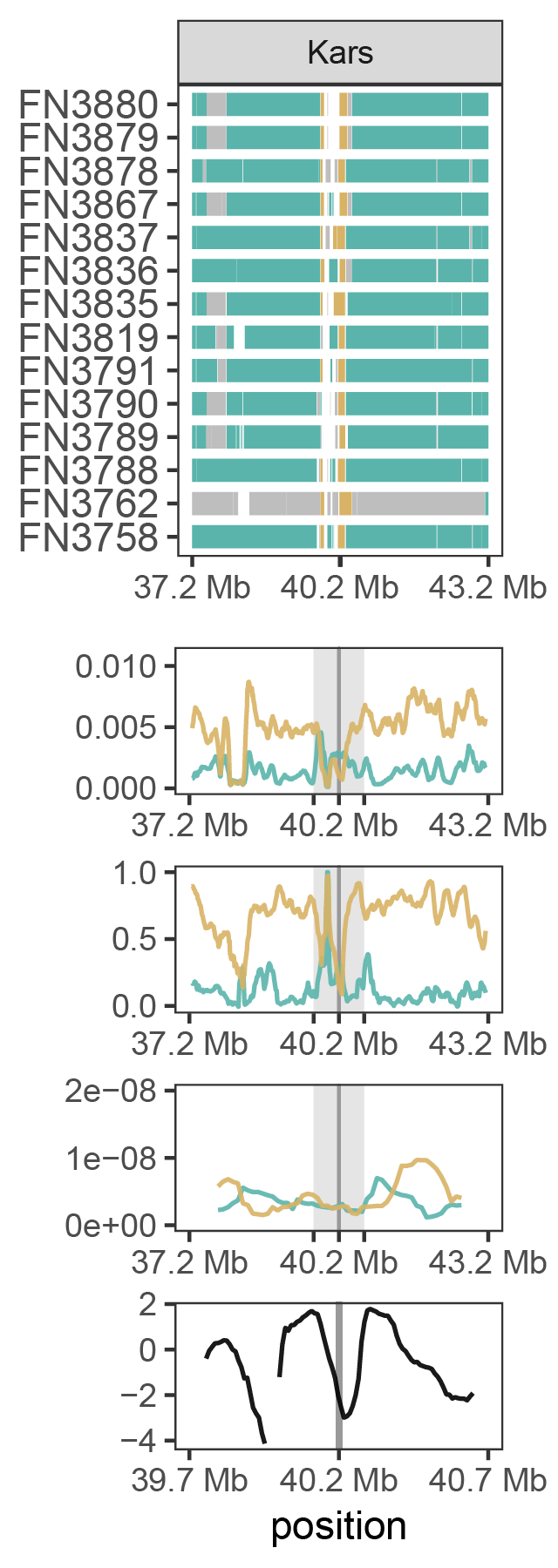

**Figures S8-S10.**

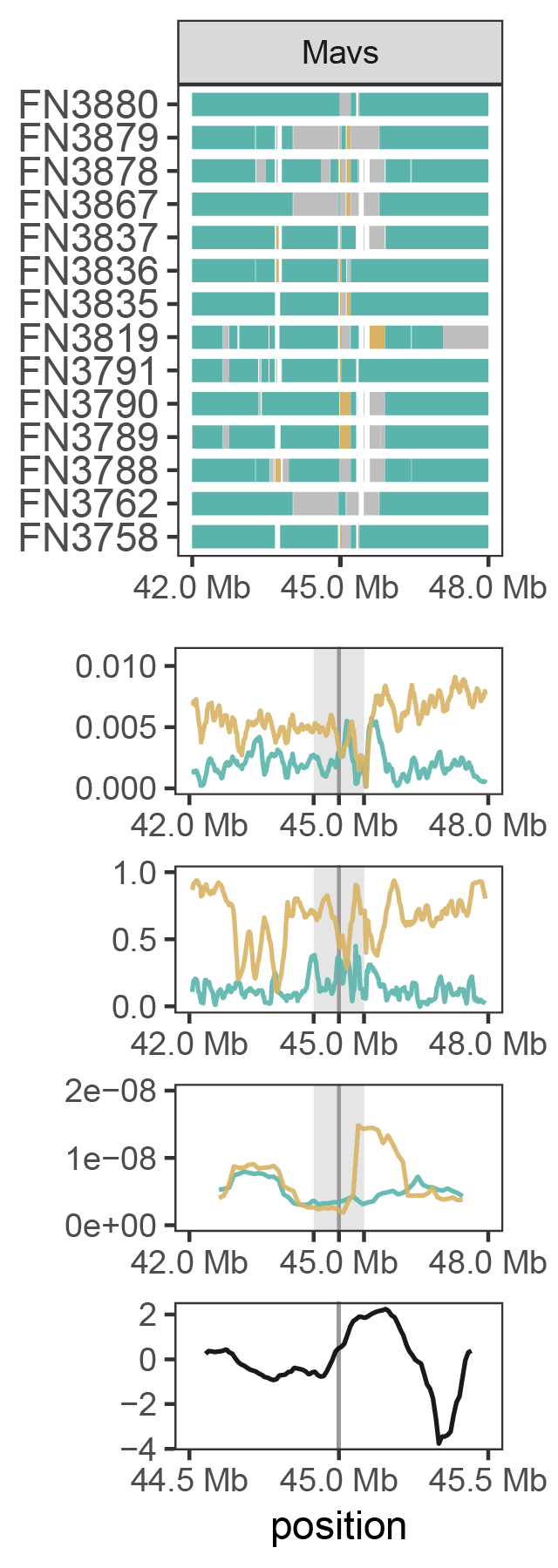

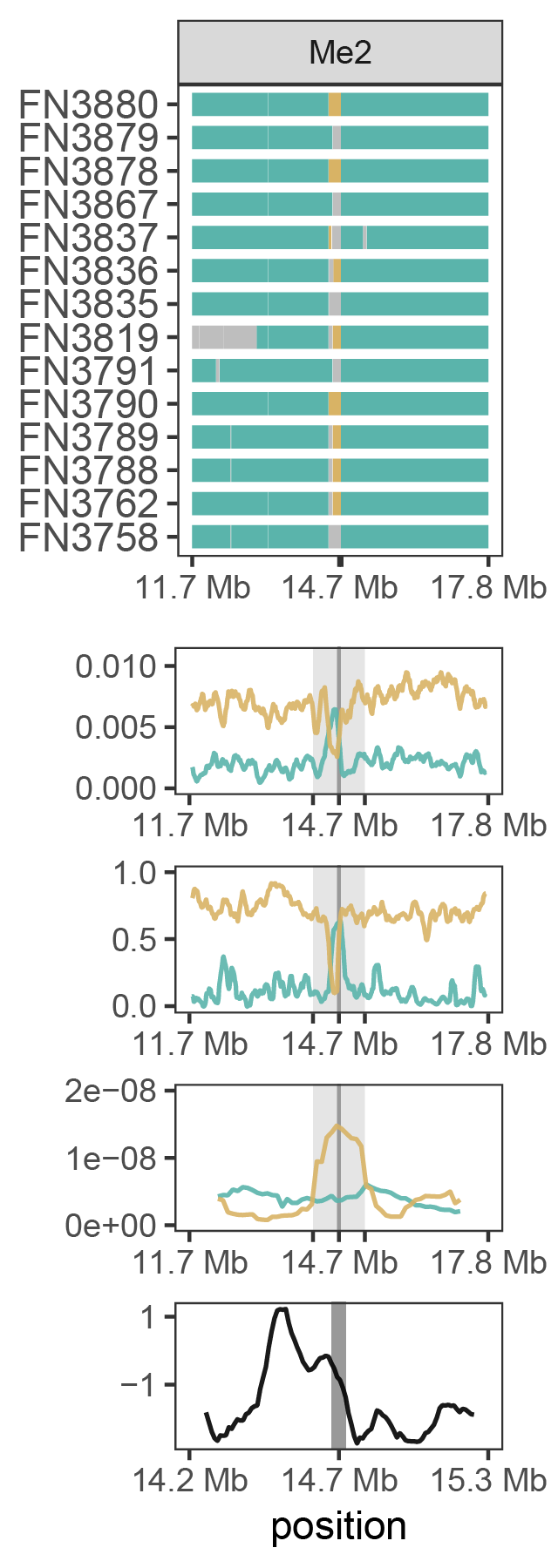

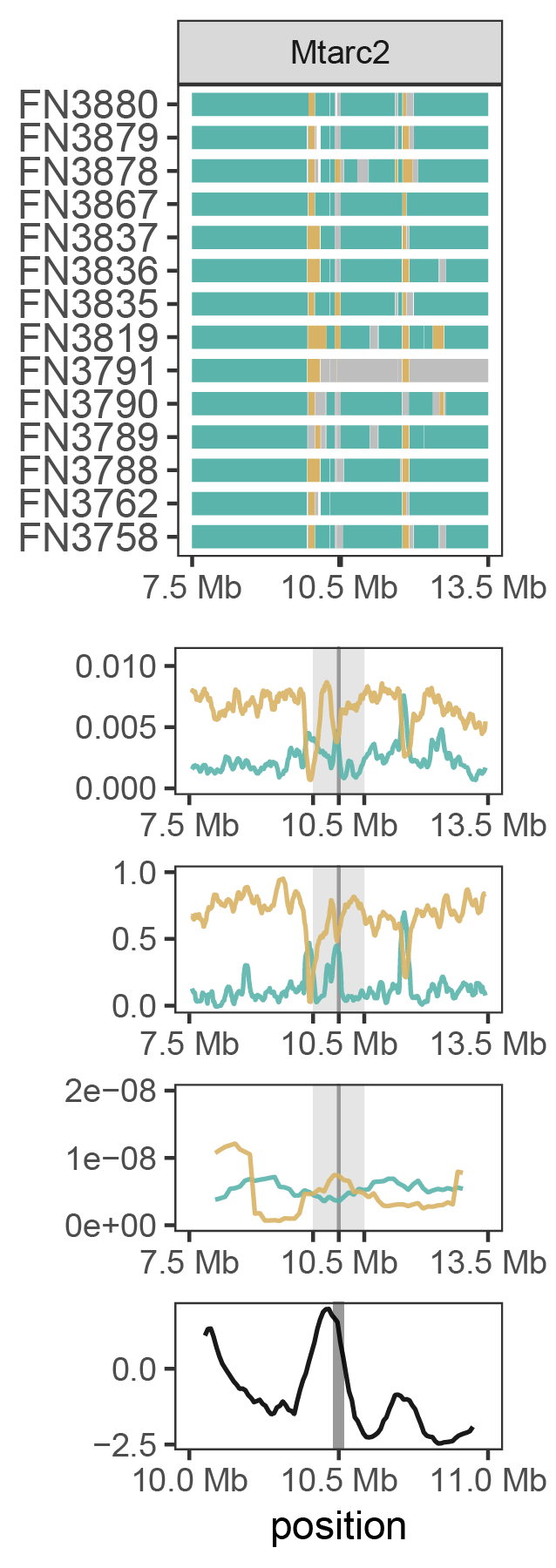

**Figures S11-S13.**

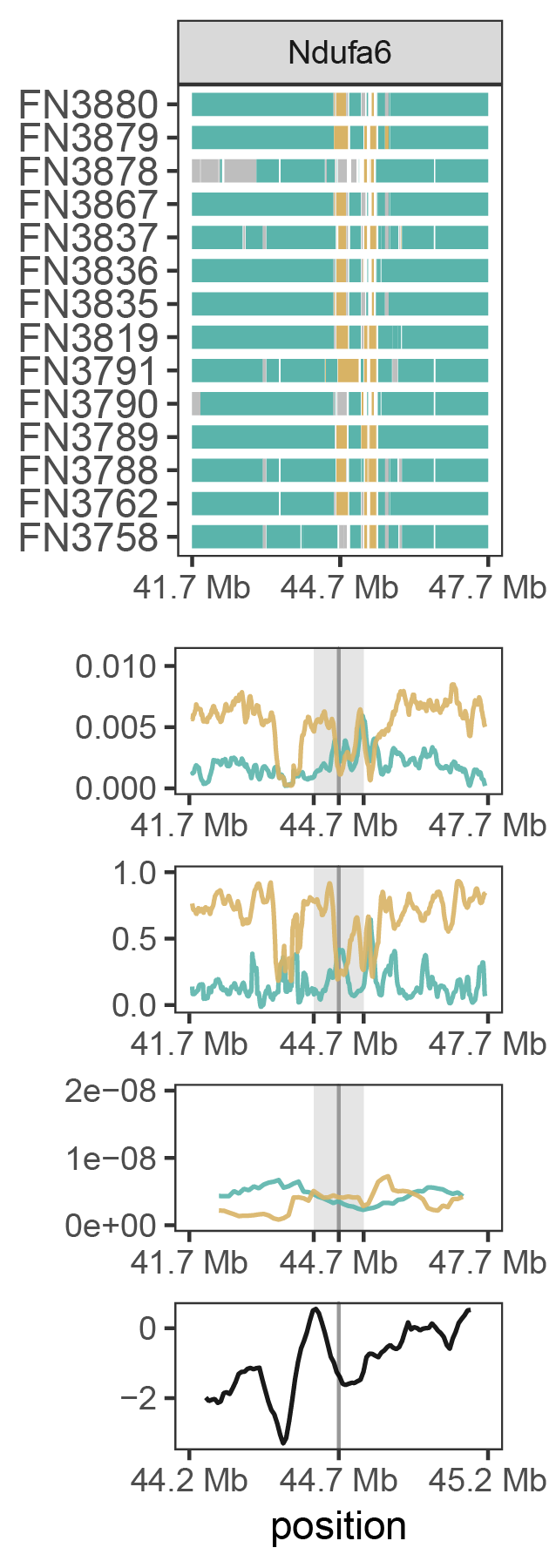

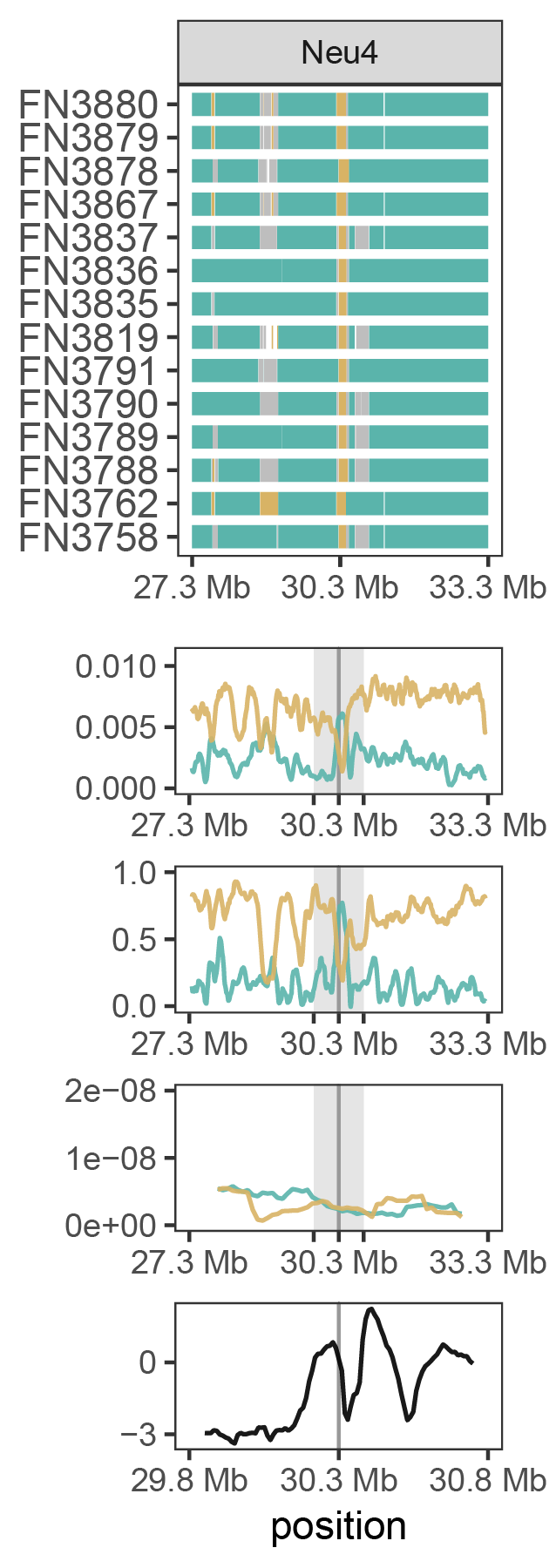

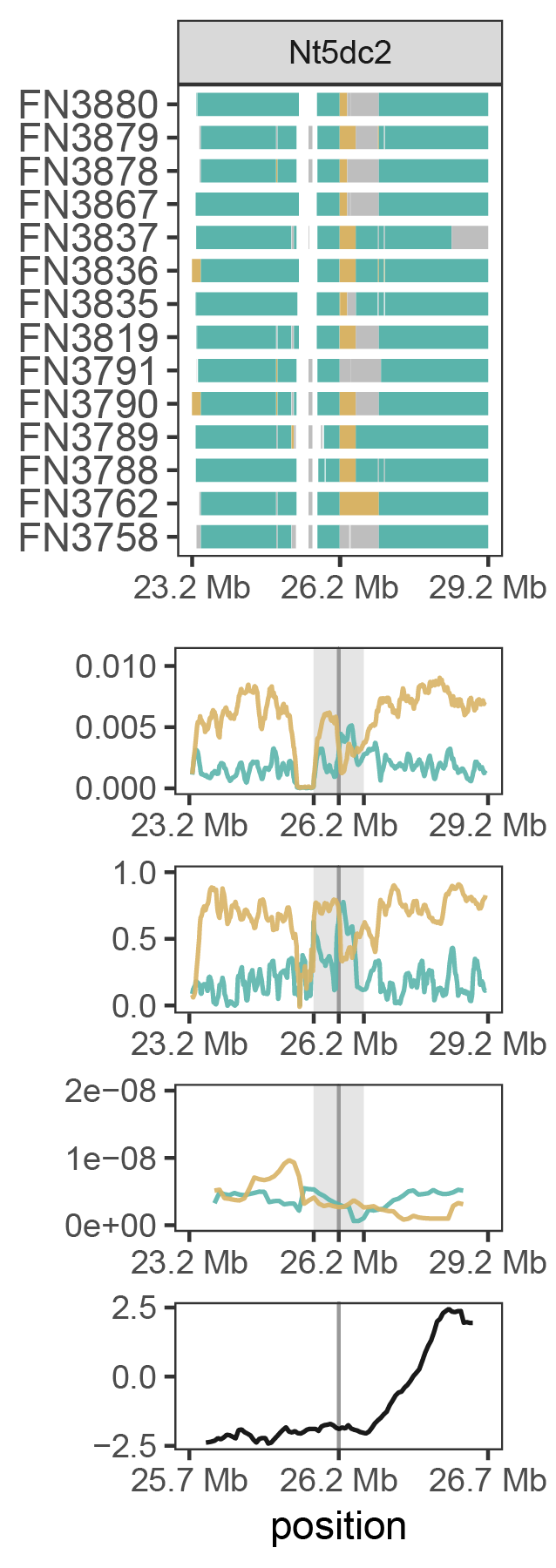

**Figures S14-S16.**

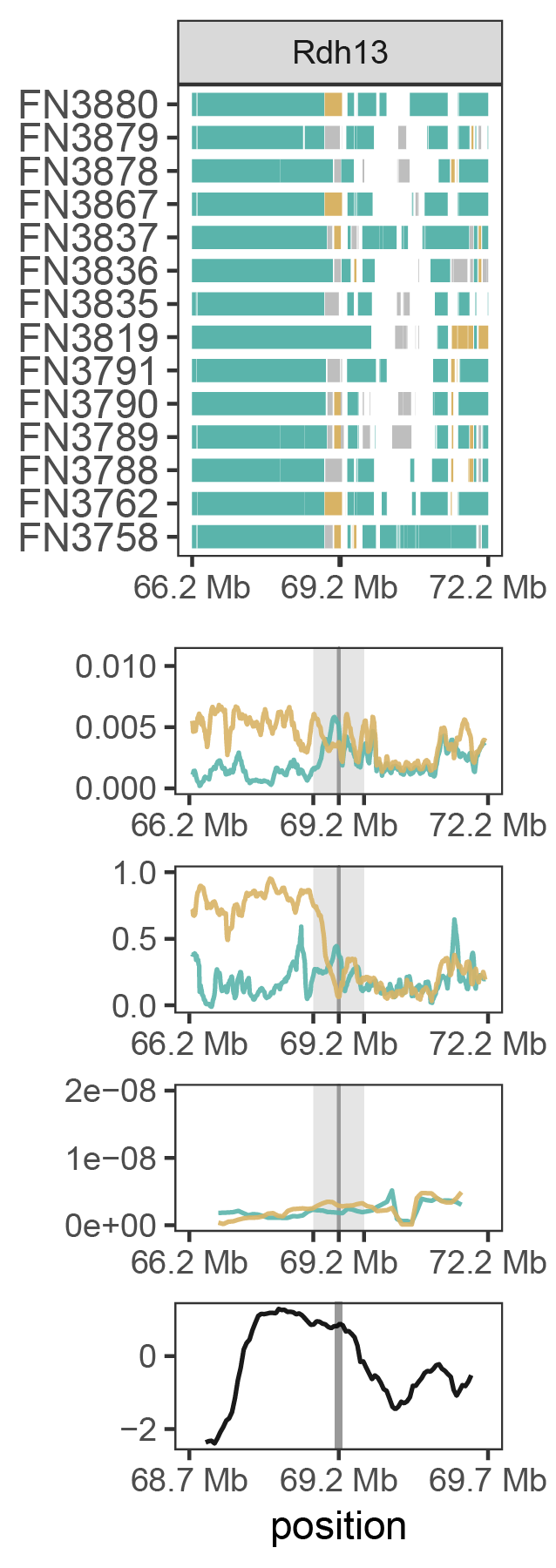

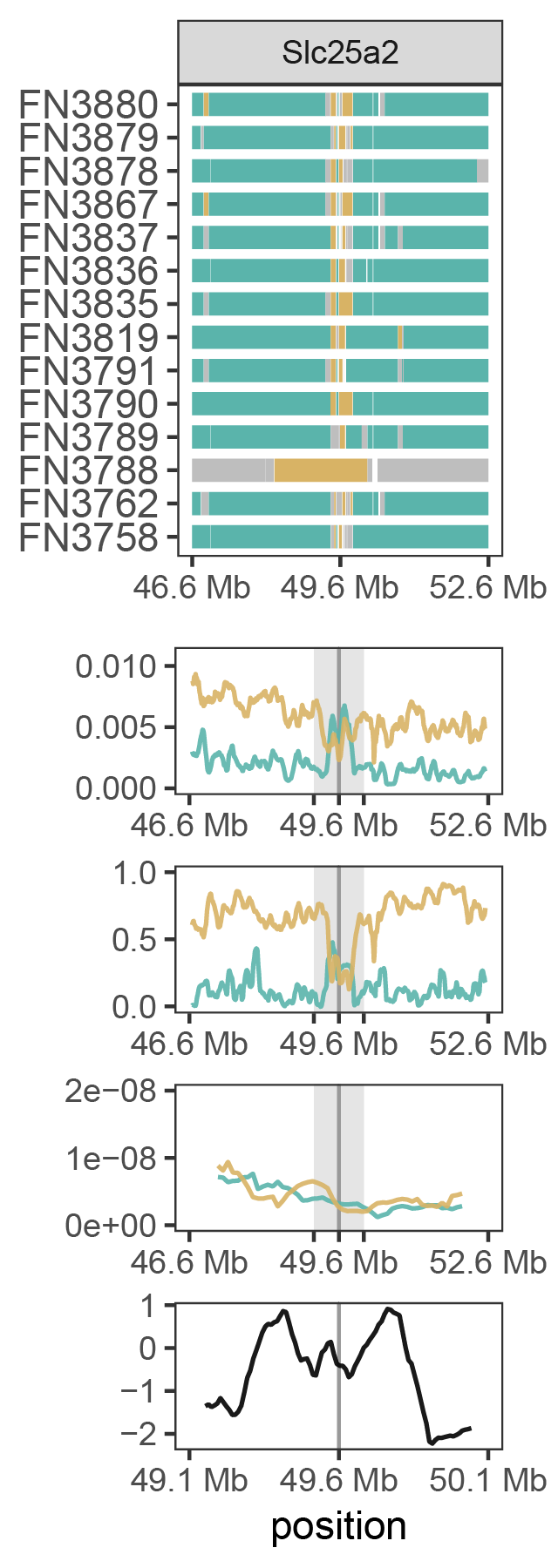

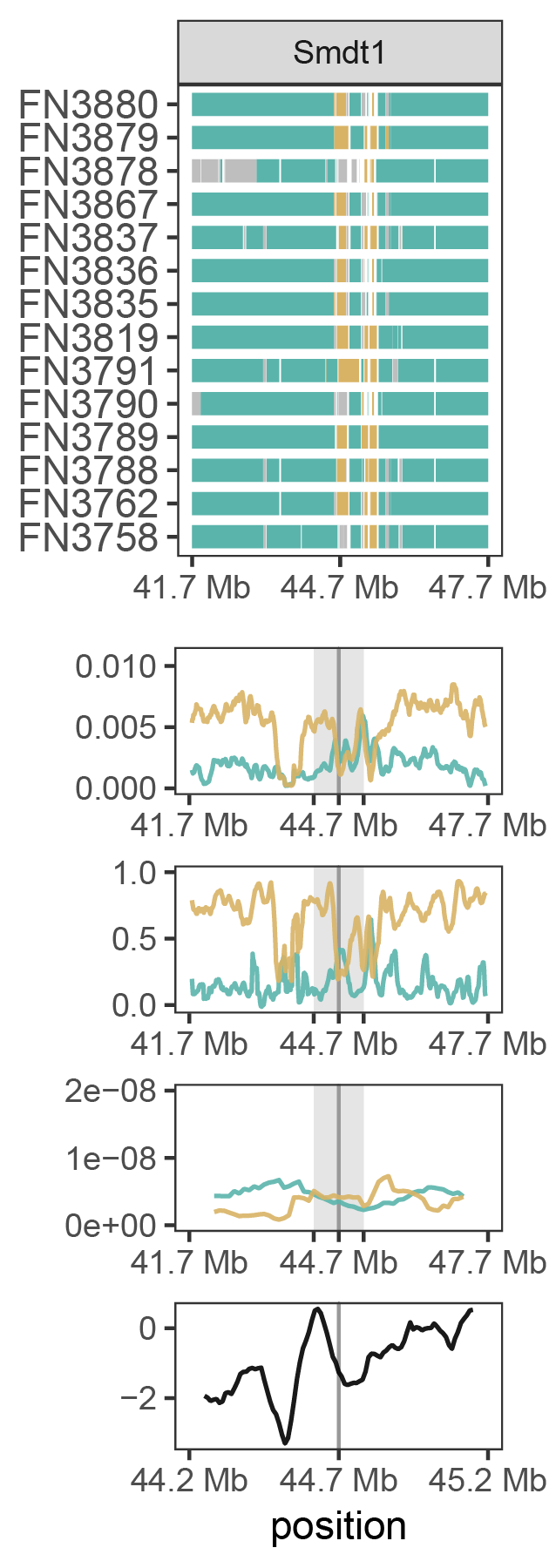

**Figures S17-S19.**

**Figures S20-37. The following pages show patterns of genomic variation in the Southeast Alaska discordant population at each N-mt gene from Table 1.** Each column represents a single gene and the surrounding region. From top to bottom, the panels show local ancestry, D_XY_, F_ST_, recombination rate, and *Tajima’s D*. The gene is centered in each panel, with its location noted by the center black tick and/or the dark grey highlighting line. The surrounding 3 Mb up- and downstream of the gene are shown for each panel, except for *Tajima’s D*, which shows the surrounding 500 kb up- and downstream of the gene. The range of genomic coordinates shown in the *Tajima’s D* panel are reflected by the light gray highlighting in the D_XY_, F_ST_, and recombination rate (r) panels. In the local ancestry panels, each individual vole is represented by a single horizontal bar. Species ancestry at each position of the chromosome is indicated by color: *C. gapperi* = teal; *C. rutilus* = gold; heterozygous = grey. Positions with less than 95% posterior probability for any species ancestry are white. In the D_XY_ and F_ST_ panels, genetic differentiation between the southeast Alaska discordant population and each reference population (comparison to *C. gapperi* = teal; comparison to *C. rutilus* = gold) is summarized in 100 kb windows, slid across the region in 10 kb intervals. Recombination rate (r) is summarized for each reference population in 1 Mb windows, slid across the region in 100 kb intervals. *Tajima’s D* is summarized for the southeast Alaska discordant population in 100 kb windows, slid across the region in 10 kb intervals.

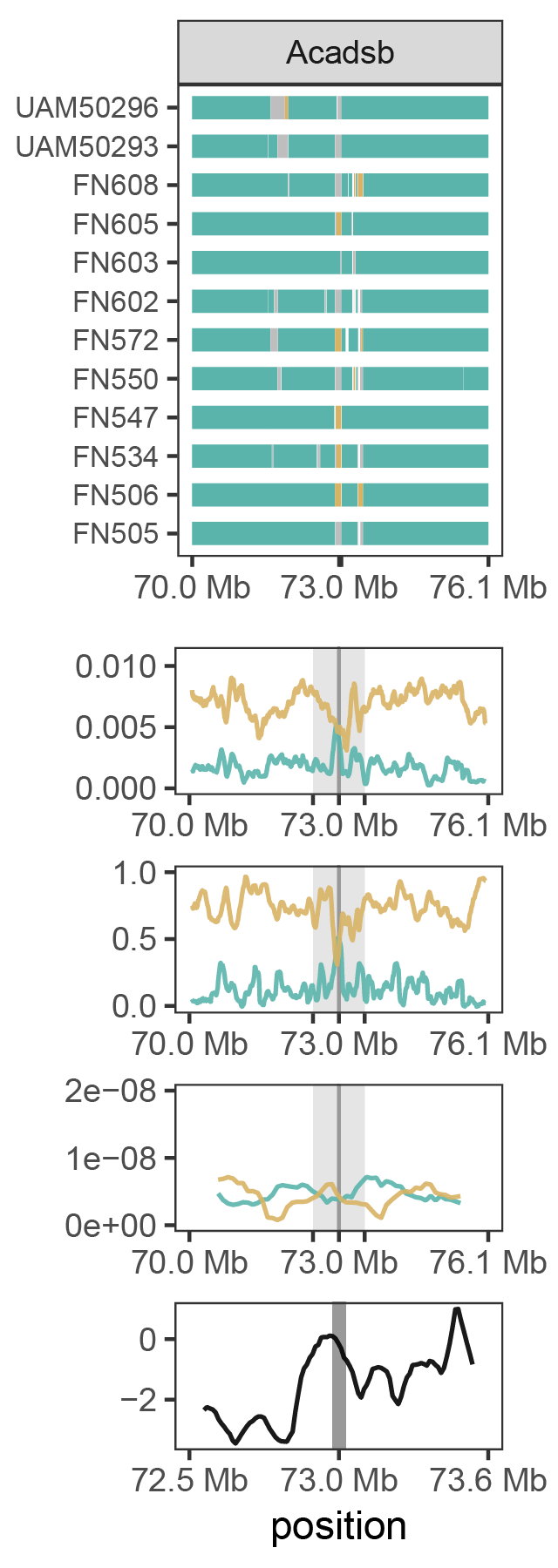

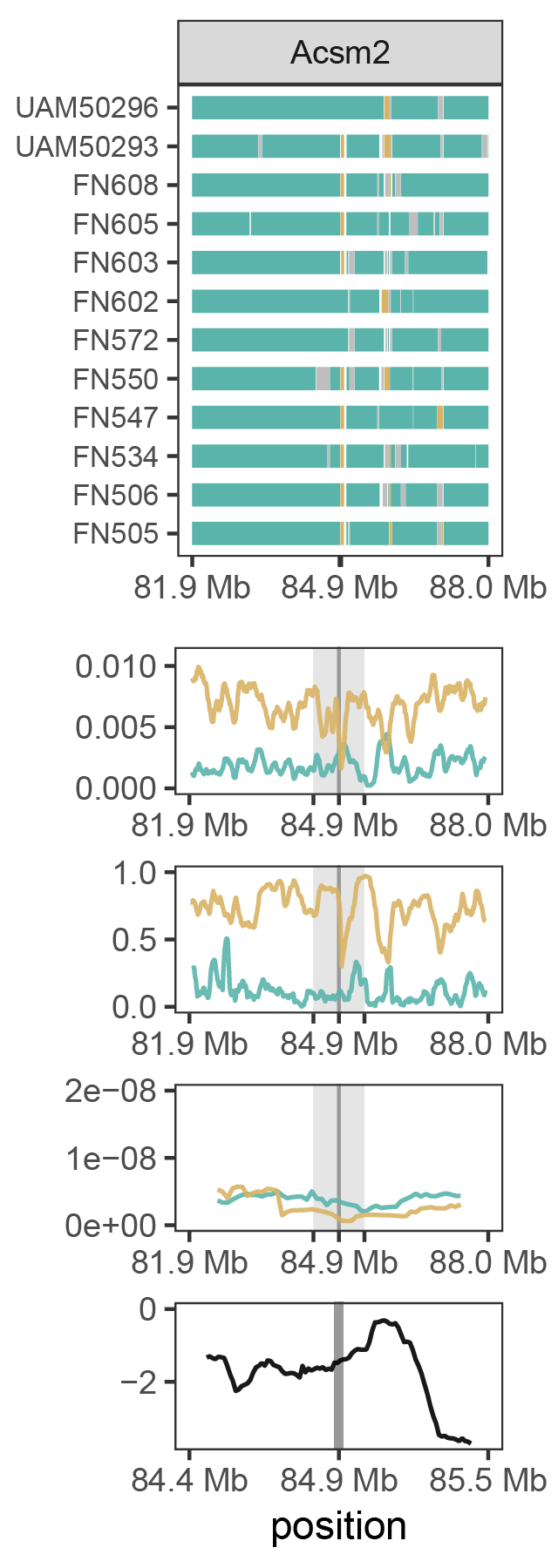

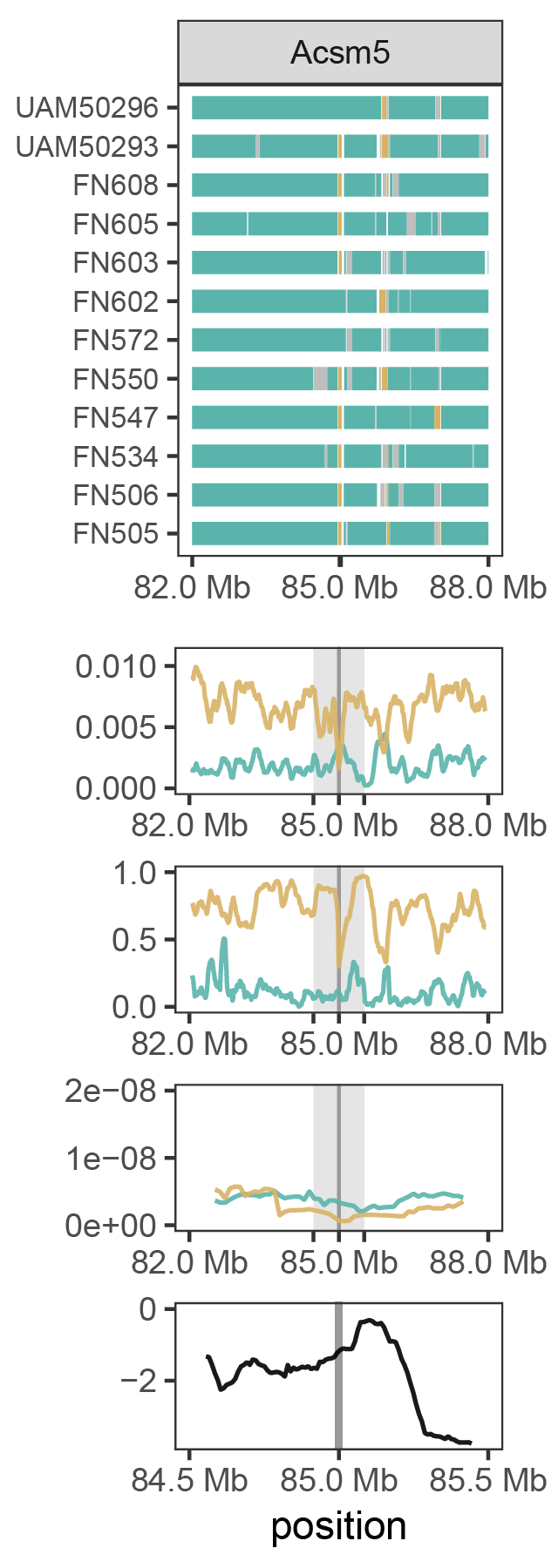

**Figures S20-S22.**

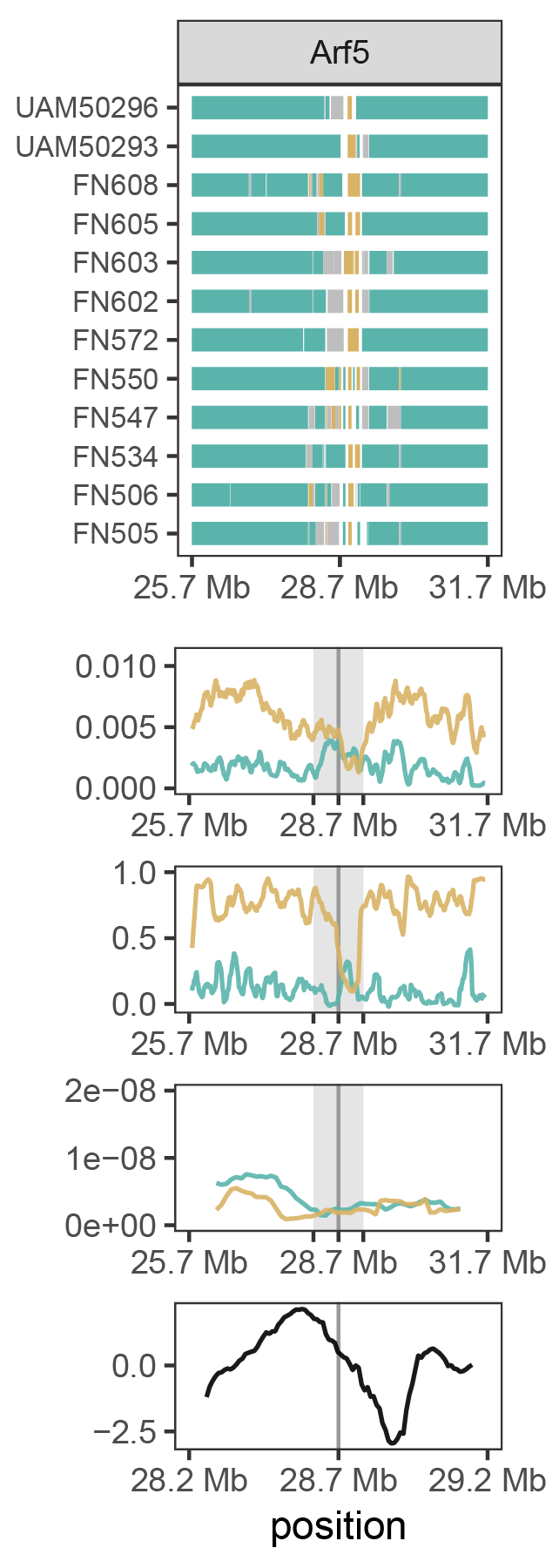

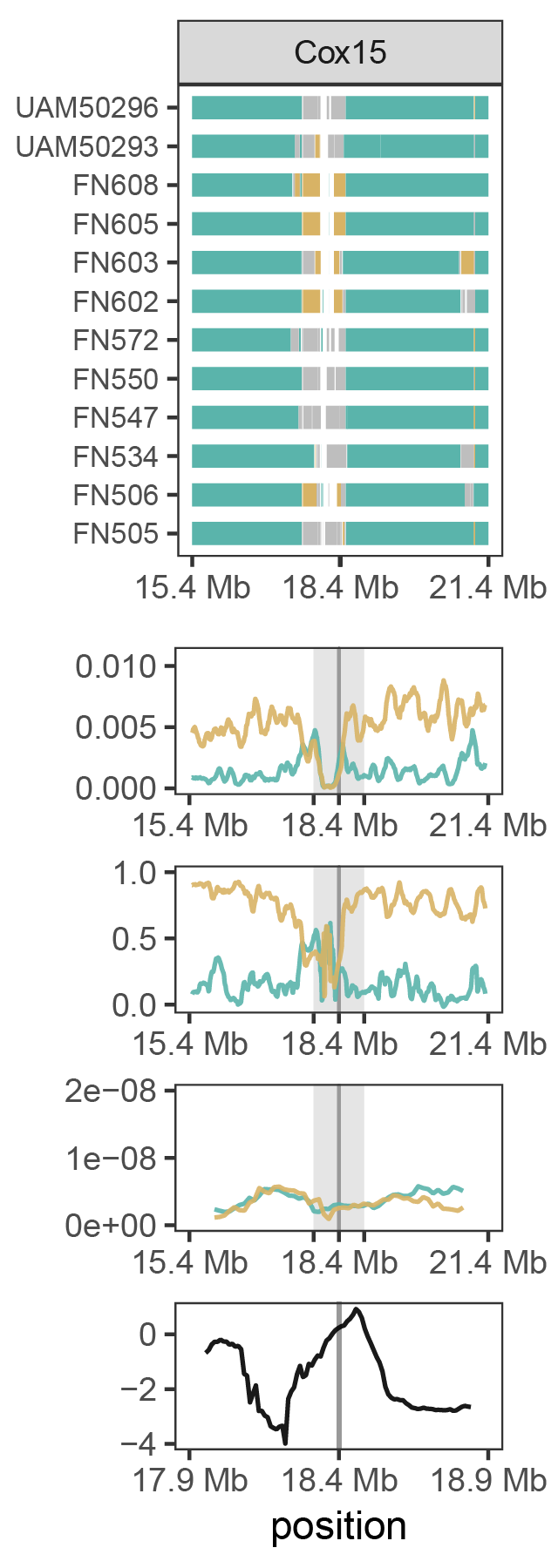

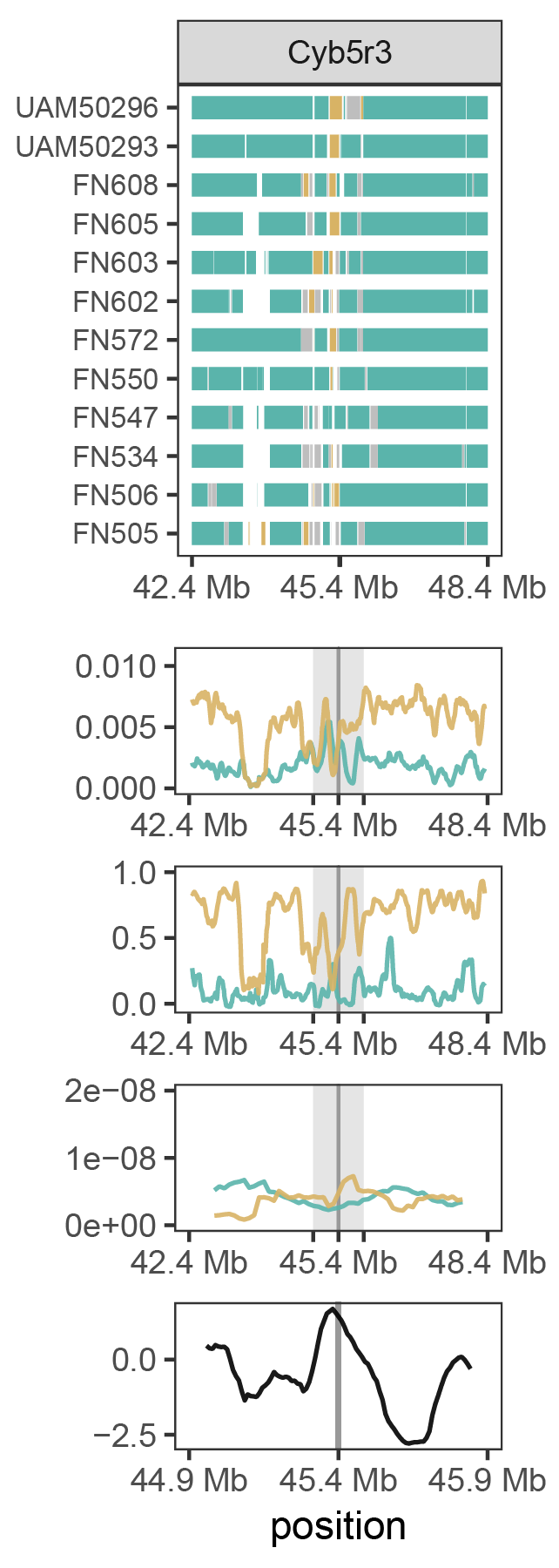

**Figures S23-S25.**

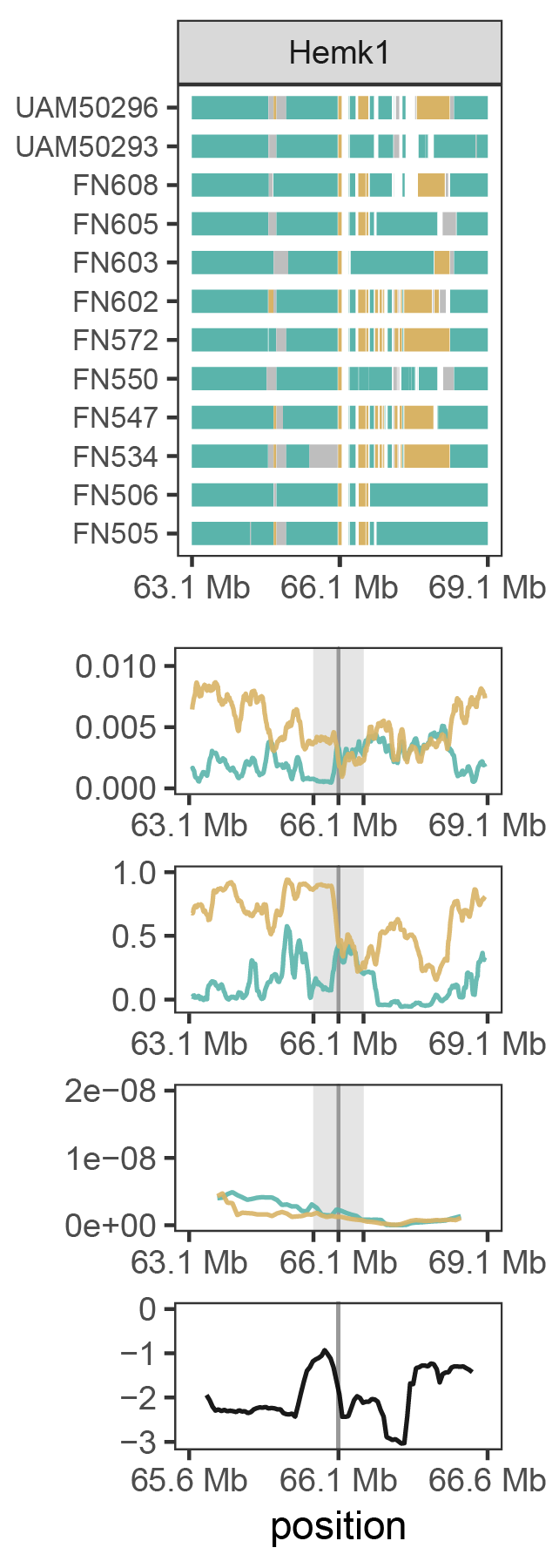

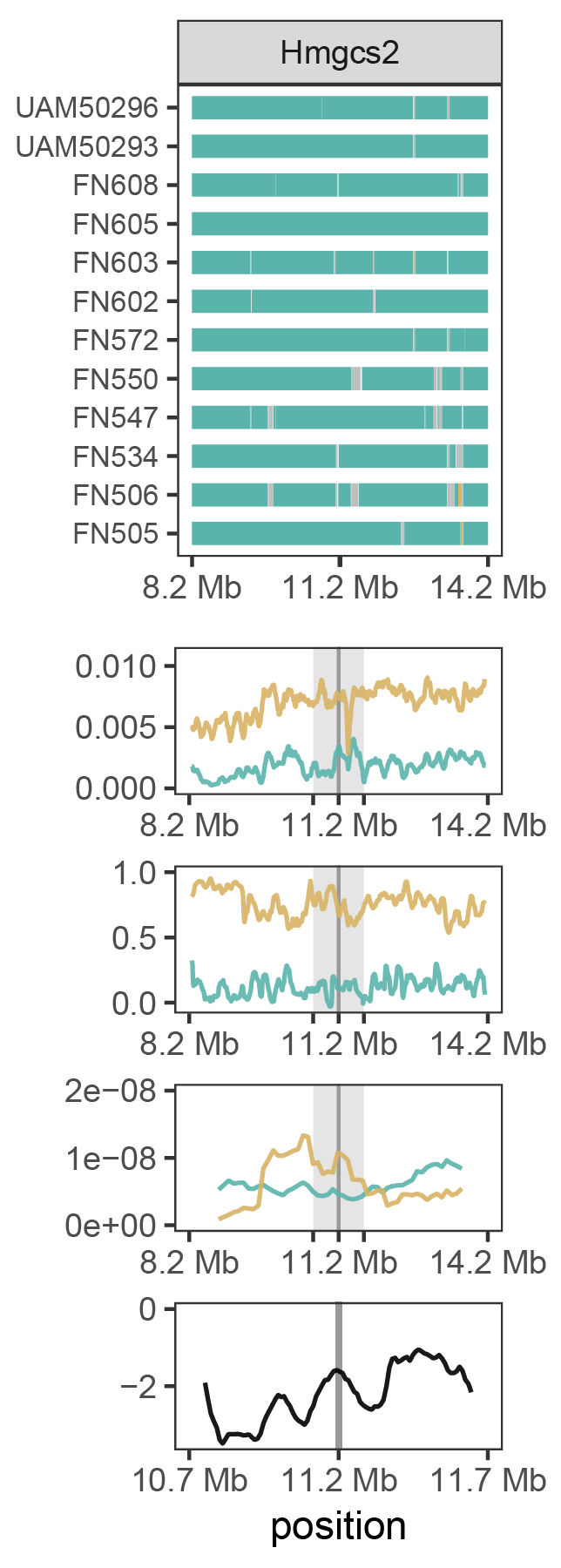

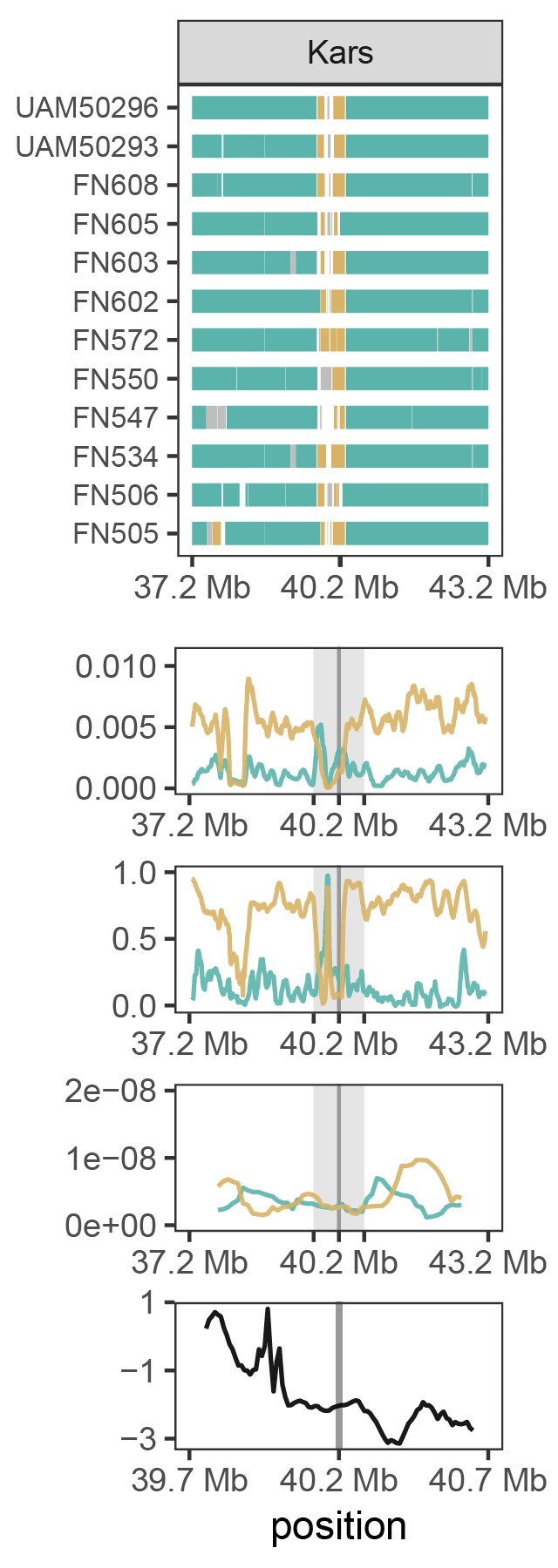
**Figures S26-S28.**

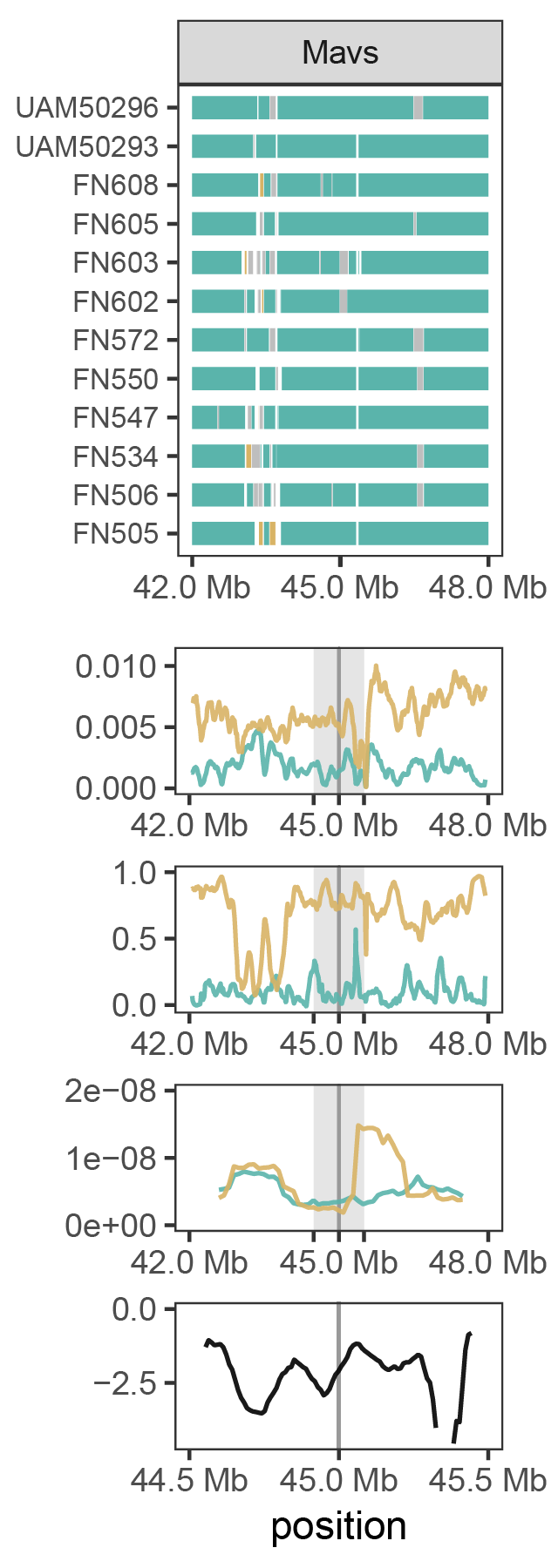

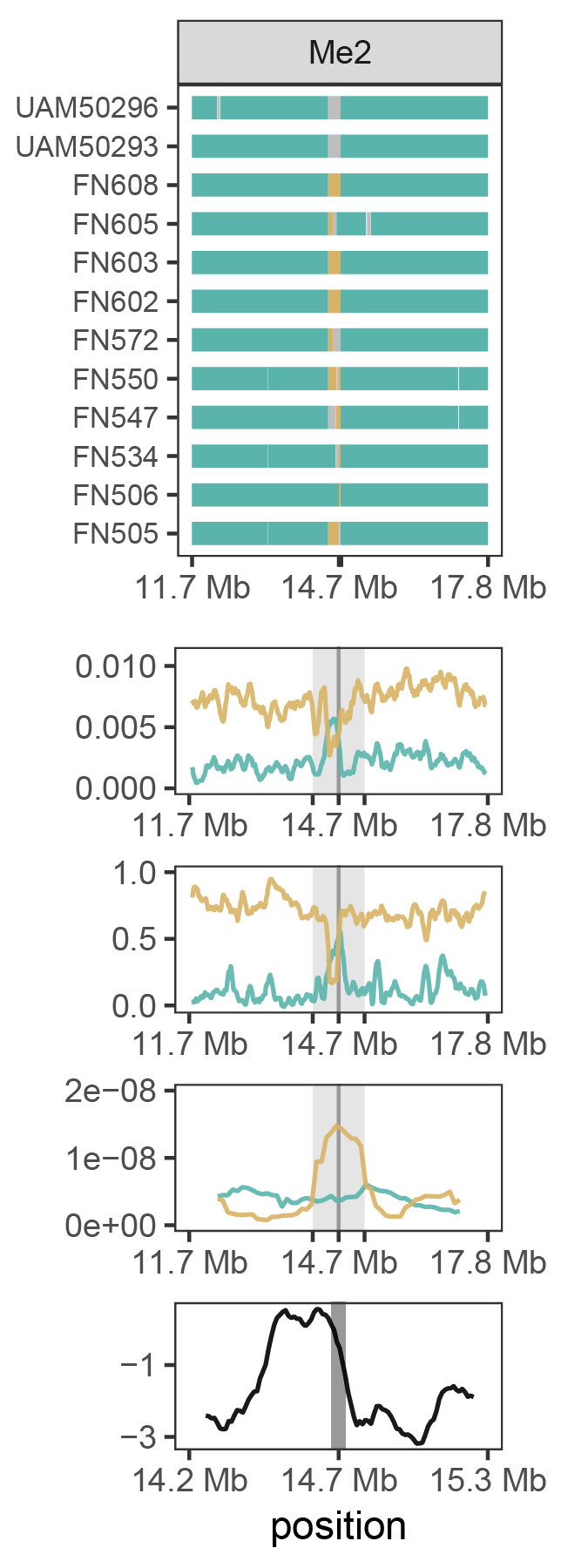

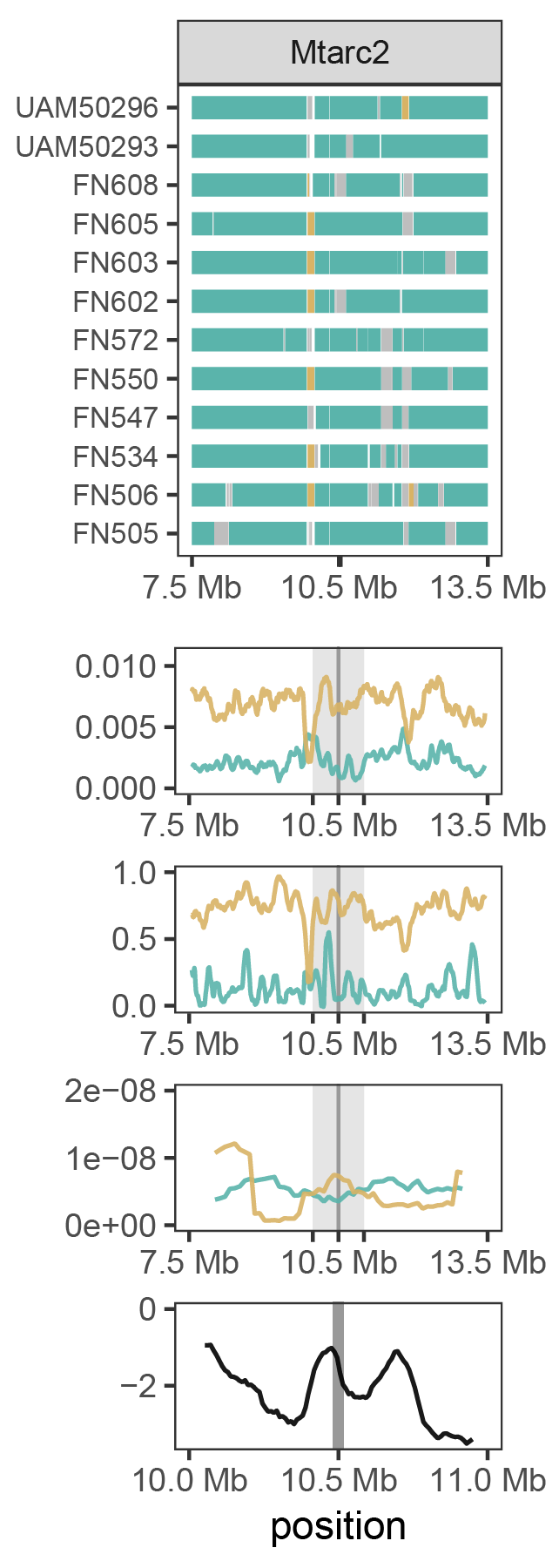

**Figures S29-S31.**

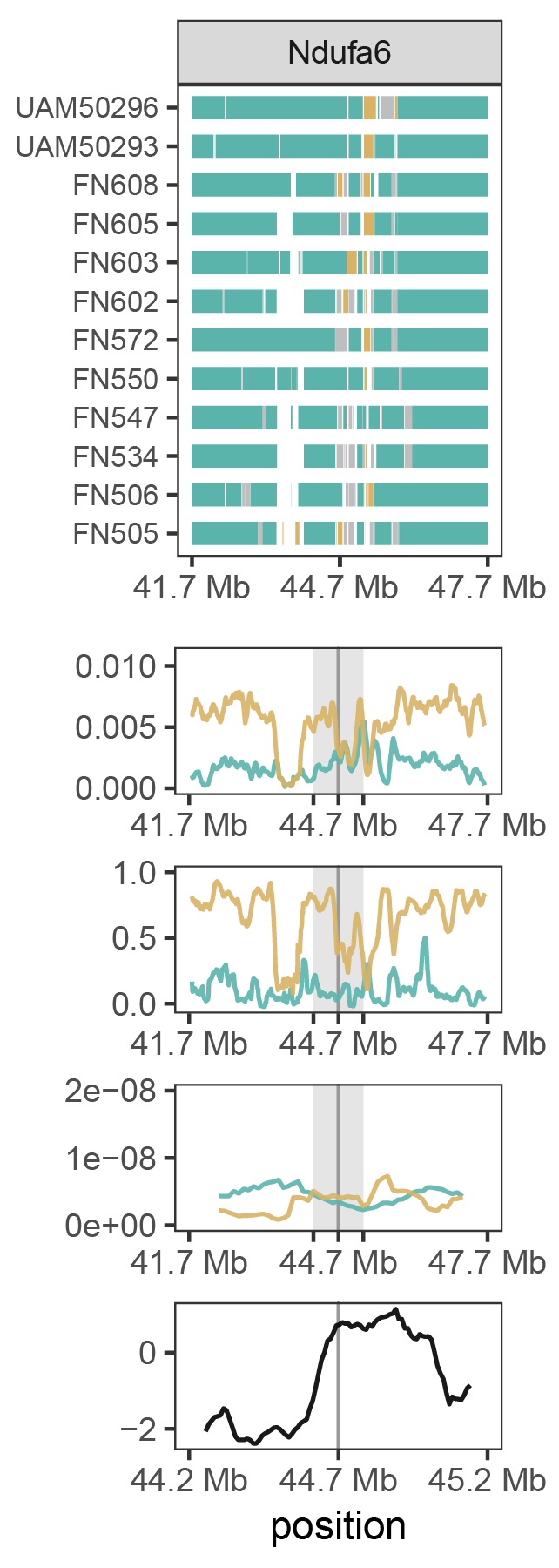

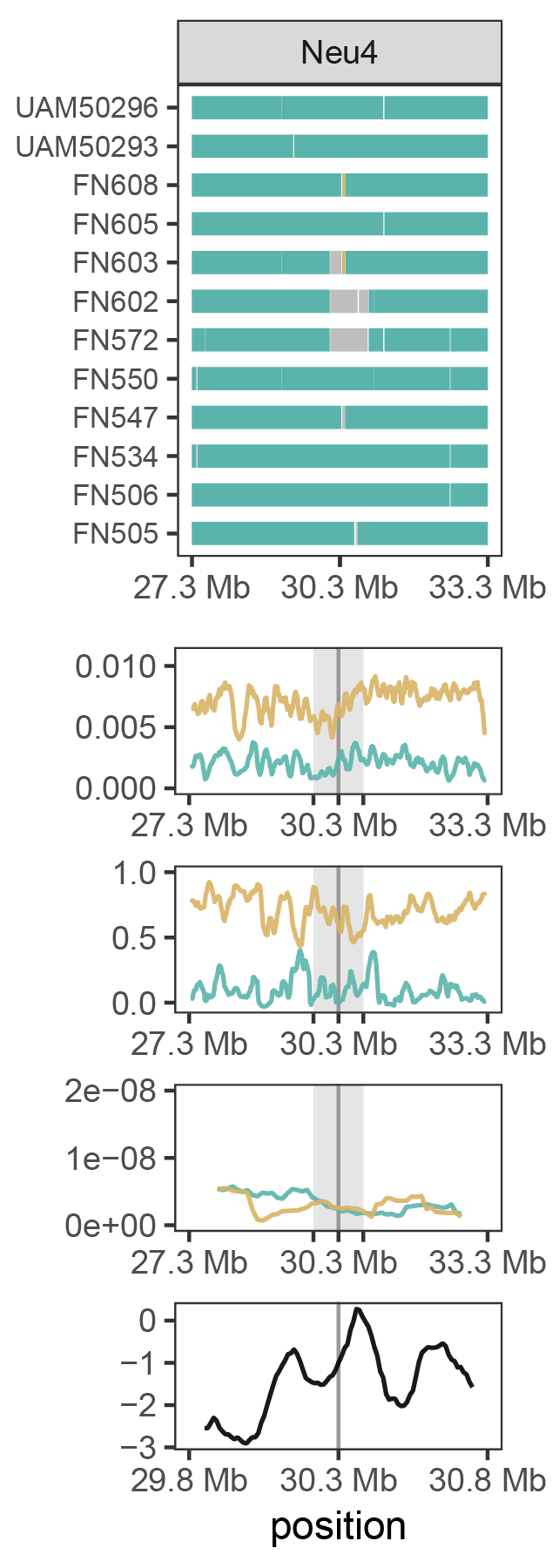

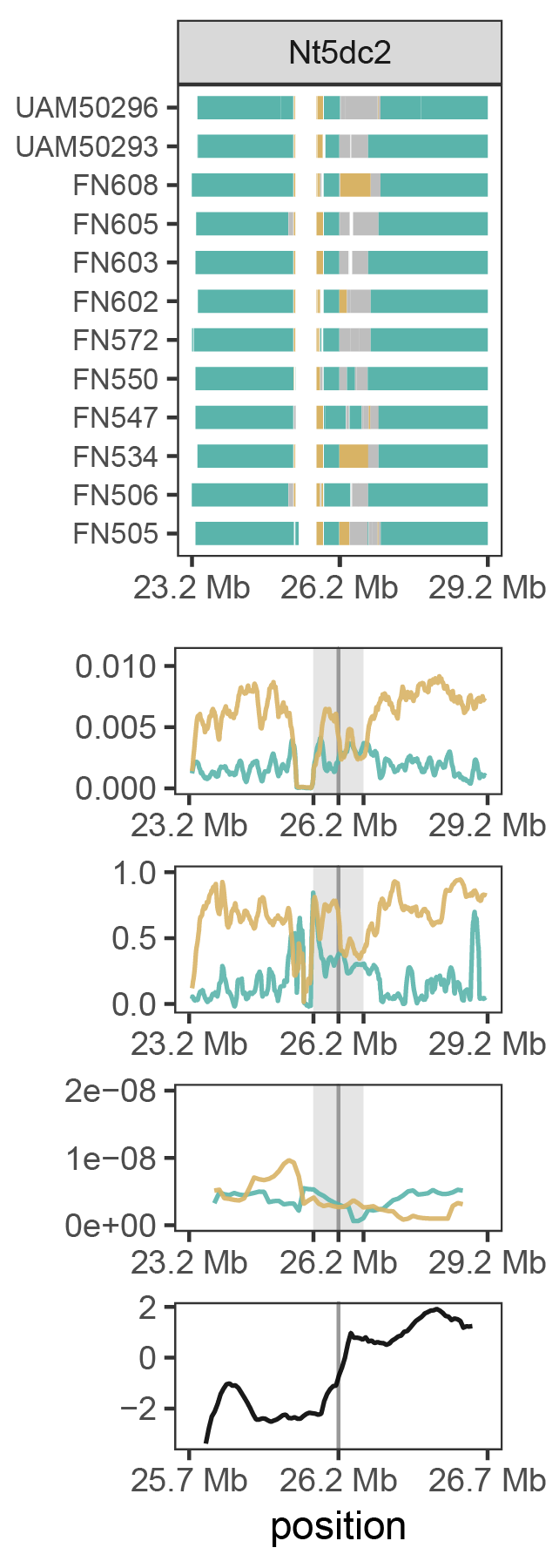

**Figures S32-S34.**

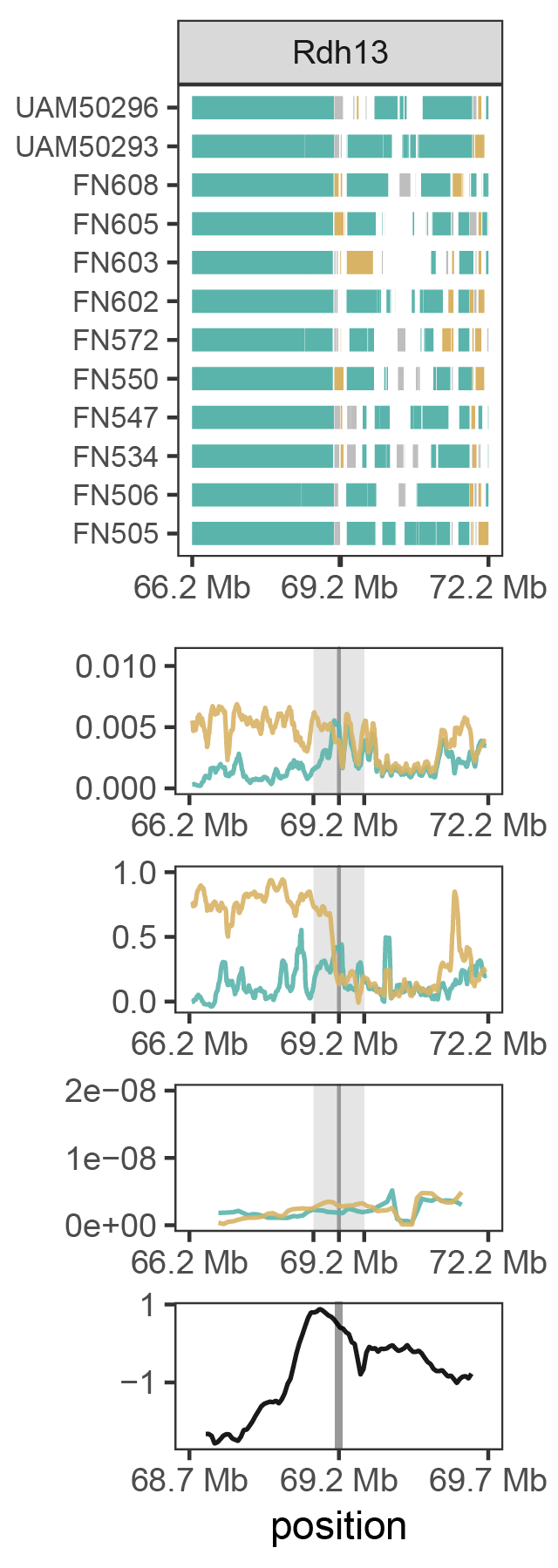

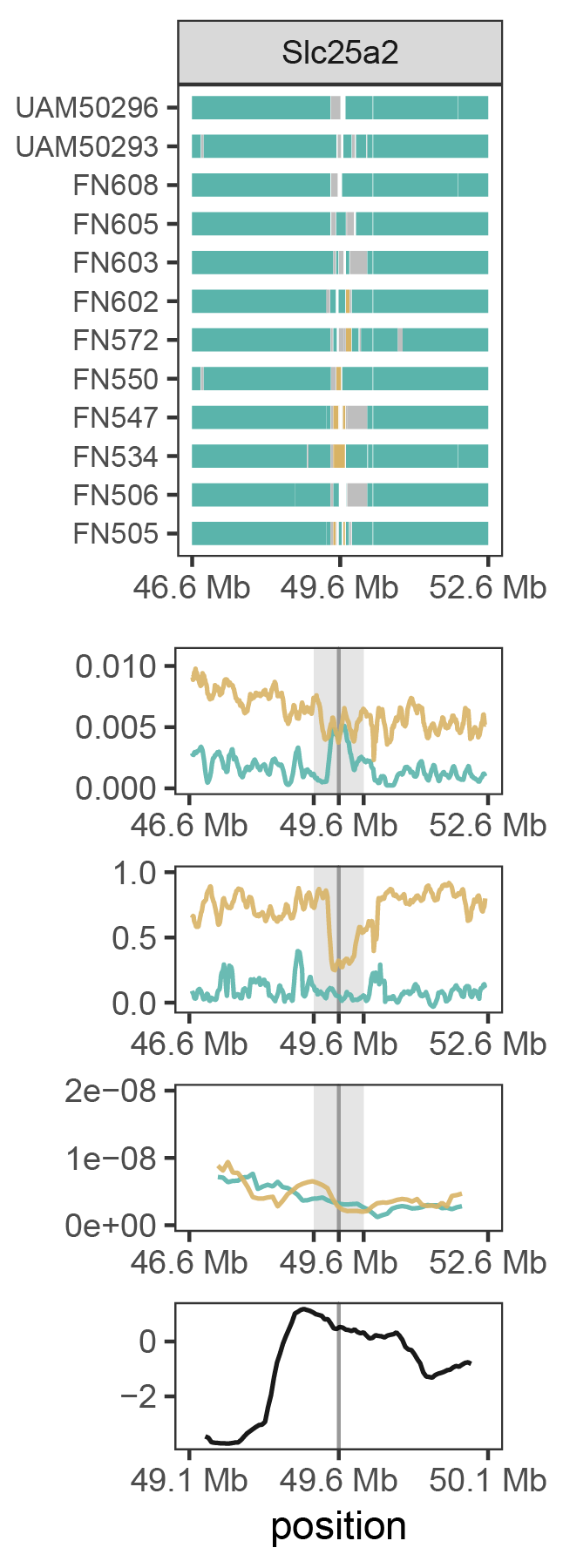

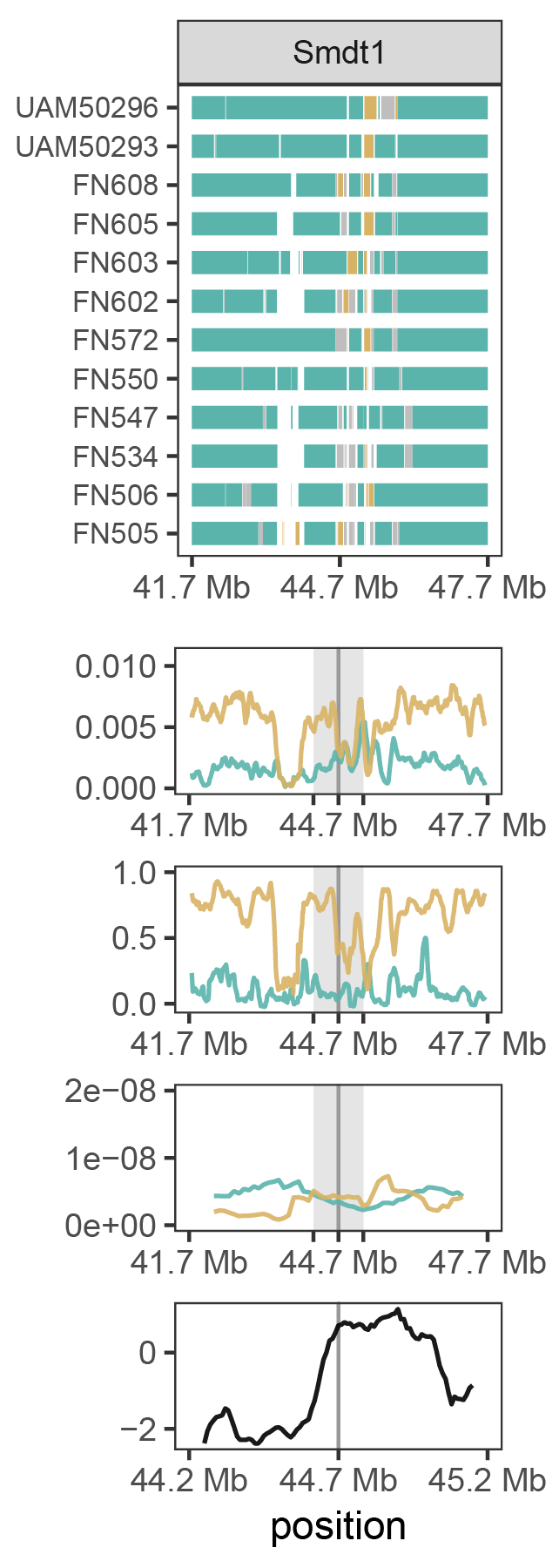

**Figures S35-S37.**
